# Evolution of oncogene amplification across 86,000 cancer cell genomes

**DOI:** 10.64898/2026.02.12.705658

**Authors:** Jake June-Koo Lee, Sohrab Salehi, Matthew A. Myers, Marc J. Williams, Melissa A. Yao, Duaa H. Al-Rawi, Jin Lee, Eric G. Sun, Kerstin Thol, Seongmin Choi, Eliyahu Havasov, Asher Preska Steinberg, Michelle Wu, Nicole Rusk, Caitlin Timmons, Cheryl Zi Jin Phua, Stephen Martis, Neeman Mohibullah, Parvathy Manoj, Esther Redin, Álvaro Quintanal-Villalonga, Pedram Razavi, Samuel Aparicio, Natasha Rekhtman, Viviane Tabar, Mark M. Awad, Helena A. Yu, Kenny Kwok Hei Yu, Andrea Ventura, Andrew McPherson, Charles M. Rudin, Sohrab P. Shah

## Abstract

High-level copy-number (CN) amplification (HLAMP) is a major mechanism of oncogene activation in human cancer. Despite progress in therapeutically targeting amplified oncogenes, the processes underlying amplicon evolution remain incompletely understood, leaving critical knowledge gaps in their etiology and mechanisms of therapeutic response. To address this, we analyzed the evolutionary trajectories of HLAMPs using single-cell whole-genome sequencing data from 86,239 cancer cells across 93 patients and 9 experimental systems. We found that cell-to-cell CN variability provides a quantifiable readout of HLAMP mechanism, clearly distinguishing extrachromosomal circular DNA (ecDNA) from intrachromosomal amplification (ICamp) through characteristic CN distributions that reflect distinct modes of segregation and correspond to clonal architecture. Notably, ICamp events frequently showed multiple amplitude peaks specific to subclones, indicating punctuated shifts in oncogene dosage through numeric or structural modulatory mechanisms with transcriptional impact. In contrast, ecDNAs exhibited broad, continuous CN distribution with extreme high-copy outliers, consistent with asymmetric segregation. The CN and structural diversity of ecDNA regions enabled systematic deconvolution of ecDNA subspecies and estimation of their per-cell abundance, revealing the history of ecDNA-mediated oncogenesis at single-nucleotide resolution. We observed ecDNA diversification through internal rearrangements across cases and, notably, convergent evolution in glioblastoma cases marked by multiple, recurrent acquisition of *EGFR*-targeting ecDNAs. Finally, single-cell genome-based identification of ecDNAs showed substantial discrepancy with bulk genome graph-based predictions and reliably distinguished actively maintained ecDNAs from historical genomic footprints after chromosomal re-integration. These findings reveal marked tissue-type specificity of ecDNAs, suggesting that ecDNA-mediated oncogenesis may depend on a permissive tissue context.

## Introduction

High-level copy-number amplification (HLAMP) is a common mechanism of oncogene activation in human cancers and often creates a targetable functional dependence. For example, anti-HER2 therapies in HER2-positive breast cancer^1^ and other cancer types^2,3^ have enabled effective precision cancer medicine in previously intractable tumors. In contrast to other primary oncogenic alterations, such as point mutations and gene fusions, HLAMPs often exhibit profound intratumoral heterogeneity. For example, *ERBB2* (encoding HER2) copy number can differ between primary and metastatic sites^4^, across regions within a tumor^5^, and among individual cells^6^. Such heterogeneity has been associated with poor clinical outcomes and therapy resistance^7^, yet its mechanistic basis remains incompletely understood. One plausible explanation is amplification via extrachromosomal circular DNA (ecDNA), which lacks centromeres and therefore can undergo unequal segregation during cell division^8^. Recent studies indicate that non-Mendelian inheritance^9^, together with unique biophysical properties of ecDNA^10^, can drive clonal evolution and lead to heterogeneous cancer cell populations^11,12^. By contrast, intrachromosomal amplification (ICamp) is expected to segregate symmetrically with chromosomes. How ICamps evolve and contribute to observed cellular heterogeneity remains unclear. Recently, therapeutic approaches exploiting selective vulnerabilities of ecDNA-containing cancer cells have been proposed^13–15^, underscoring the clinical importance of accurately distinguishing HLAMP mechanisms and characterizing their evolutionary dynamics.

To map the mechanisms underlying HLAMP etiology, evolution and their intratumoral heterogeneity in clinical tumor samples, we analyzed a large-scale single-cell whole-genome sequencing (scWGS) dataset consisting of >86,000 single cells genomes derived from >100 samples across a range of solid cancer types and experimental models. This enabled both direct quantification of cell-to-cell copy number (CN) variability^16,17^ and identification of structural variant (SV)-resolved segment boundaries at nucleotide resolution at HLAMP loci. From these data, we developed a probabilistic mixture model, eicicle, informed by evolutionary simulations to distinguish between ICamp and ecDNA based on single-cell CN distributions. We found that ICamps diversify at a clone level pervasively through two mechanisms: numeric modulation whereby the copy number architecture is maintained, but arm-level or chromosomal-level aneuploidies increase or decrease copy number in a clone-specific manner; and structural modulation whereby SV-driven mutational processes drive subclonal diversification. In ecDNAs we observed cases harboring multiple ecDNA species, and developed ECADeMix, a mixed-integer linear programming framework, to decompose ecDNA heteroplasmy at single-cell resolution through simultaneous inference of ecDNA structures and cellular prevalence reflective of evolutionary histories. Finally, our scWGS-based approach underscores limitations of bulk genome graph-based ecDNA predictions and reveals notable tissue-type specificity^18^ of ecDNAs, informing patient selection for emerging ecDNA-directed therapeutics.

### Single-cell copy number variance distinguishes ecDNA and ICamp

We analyzed direct library preparation (DLP+)-based scWGS data from 102 individuals (patients and experimental models), including 71 published^17,19–21^ and 31 newly generated cases (**Fig. 1a, top** and **Supplementary Table 1**). These included clinical tumor samples and patient-derived xenografts (PDXs) from 93 patients across four cancer types: high-grade serous ovarian cancer (HGSOC; n=61), invasive breast cancer including hormone receptor-negative (HR− BC; n=10; one with *ERBB2* amplification) and -positive (HR+ BC; n=5), lung cancer including small-cell (SCLC; n=8) and non-small cell (NSCLC; n=5; all with *EGFR* mutation), and glioblastoma multiforme (GBM; n=4). Although most clinical tumors were obtained prior to systemic therapy, the cohort included several pretreated tumors, including 4 HR+ BCs and 2 EGFR inhibitor-resistant, *EGFR*-mutated NSCLCs (**Supplementary Table 2**). The cohort also included many PDX samples, which were often collected after chemotherapy exposure. In addition, we studied 9 experimental models, including cell lines (n=8) and a mouse model of liver cancer harboring engineered *Mdm2*-containing ecDNA^22^ (n=1). A total of 311 scWGS libraries generated from these cases (**Supplementary Table 3**) were assessed for quality and jointly processed with matched normal bulk WGS data for somatic variant calling, as described previously^20^. After removing normal cells and low-quality cells, the dataset consisted of 86,239 high-quality cancer cells (**Fig. 1b**). To systematically analyze the generative and evolutionary mechanisms of HLAMP^23^, we integrated somatic SV breakpoints detected at the pseudobulk level with genome-wide CN bins to generate SV-aware single-cell CN profiles (**Methods**). Using pairwise CN distance between single cells, we identified subclones and reconstructed their evolutionary relationships (**Methods**). Across the 102 cases, we identified 507 HLAMP regions with i) CN at least 3 times the chromosome arm-level baseline in one or more subclones and ii) a pseudobulk CN of at least 6. These HLAMP regions typically included *bona fide* oncogenes known to be activated by HLAMP, such as *MYC*, *CCNE1*, *EGFR*, and *MDM2* (**Fig. 1c**), in addition to loci devoid of known oncogenic drivers, often amplified in conjunction with other oncogene-containing regions. For each HLAMP region (comprising one or more contiguous SV-aware segments), we evaluated CN distributions across cells, determined generative mechanisms from SV features at their boundaries, and analyzed clone-specific features to delineate mechanisms driving subclonal diversification (**Fig. 1a, bottom**).

**Fig. 1.**
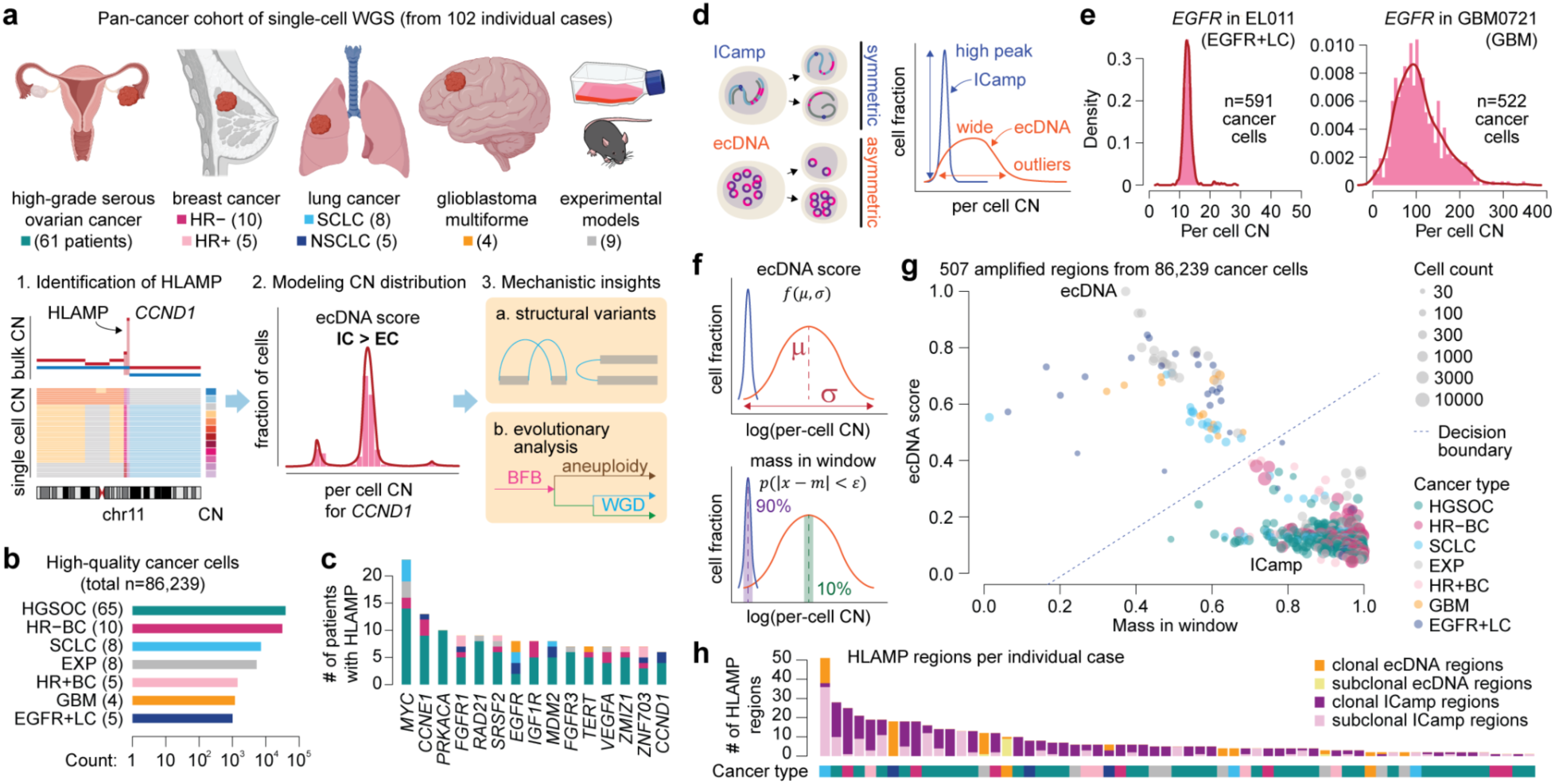
Cell-to-cell copy-number variance in scWGS differentiates ecDNA-mediated amplifications from intrachromosomal amplifications. **a**, Cancer types and case counts (top) and schematic illustration of study design (bottom). Organ and model illustrations are from BioRender. **b,** Number of high-quality cancer cells analyzed (x-axis, log-scale) by tumor type (y-axis). **c,** Frequently amplified oncogenes and their case counts. **d,** Inheritance patterns for intrachromosomal amplification (ICamp) versus extrachromosomal circular DNA (ecDNA; left) and expected CN distributions across cells (right). **e,** Representative examples of ICamp and ecDNA, from a patient with EGFR-mutated lung adenocarcinoma (left) and glioblastoma (right). **f,** Conceptual illustration of two metrics used to classify HLAMP events. **g,** Classification of 507 HLAMP regions from 86,239 cancer cells based on single-cell CN distribution. **h,** Fraction of clonal and subclonal HLAMP regions by mechanism, across cases.

First, we performed *in silico* simulation of single-cell CN distributions accounting for the mode of inheritance of oncogene amplifications and evolutionary constraints (**Methods**)^9,12,24,25^. These simulations showed that conditioned on symmetric cell division, as expected for ICamps, CN distributions consist of prominent, narrow peaks reflecting cell populations that share one or a discrete set of CN values (**Fig. 1d)**. By contrast, conditioned on asymmetric cell divisions, as expected for ecDNA^12^, broad, heavy-tail distribution with extreme outliers were observed, reflecting CN variance across the cell population. Modulating key parameters further simulated fitness consequences on ecDNA CN distributions: stronger positive selection (e.g., higher potency of oncogene) shifted the modal CN to the right, while greater cell-death penalties (e.g., DNA damage responses at extreme CN) shifted it to the left (**Extended Data Fig. 1** and **Methods**). Consistent with these simulations, we observed two distinct and quantitatively distinguishable classes of single-cell CN distributions across HLAMP regions in our cohort. For example, in the *EGFR*-mutated lung cancer case EL011, the *EGFR* allele harboring the E746-A750 microdeletion was highly amplified and exhibited a narrow, high-CN peak from 591 cancer cells, consistent with ICamp (**Fig. 1e, left** and **Extended Data Fig. 2a**). In contrast, the surgically resected glioblastoma GBM0721 showed a broad, heavy-tail distribution of wild-type *EGFR* CN across 522 cancer cells, consistent with ecDNA (**Fig. 1e, right** and **Extended Data Fig. 2b**). Motivated by these patterns, we developed a machine learning-based classifier, eicicle, to distinguish ecDNA from ICamp (**Fig. 1f**). Eicicle includes two metrics: *ecDNA score* which captures distribution width, and *mass-in-window* which quantifies the fraction of cells near the modal CN (high in ICamp, low in ecDNA). These metrics are derived from a probabilistic mixture model that conditionally models the two distinctive distributions (**Methods**). The 507 HLAMP regions by these metrics were bimodally distributed (**Fig. 1g**). We performed a discriminant analysis to infer a decision boundary, designating 15 HLAMP regions with high ecDNA scores and low mass-in-window (left upper corner) as representative ecDNA and 6 regions with low ecDNA scores and high mass-in-window (right lower corner) as representative ICamp (**Extended Data Fig. 3**). This boundary classified 72 HLAMP regions from 11 cases as ecDNA and 435 regions from 55 cases as ICamp. We then orthogonally validated classifier predictions across all available experimental models (5 ecDNA-positive and 3 ecDNA-negative by eicicle; **Extended Data Fig. 4a-c**) using metaphase DNA FISH. In all models classified as ecDNA, we observed dispersed extrachromosomal fluorescent signals for the amplified oncogenes, consistent with ecDNA. In models classified as ICamp, we observed strong intrachromosomal clustered signals. Notably, HLAMP regions classified as ecDNA were largely from GBM, SCLC, and *EGFR*-mutated NSCLC. Despite numerous HLAMP events in triple-negative breast cancers and ovarian cancers, none exhibited a characteristic single-cell CN distribution for ecDNA. Instead, they showed prominent peaks typical for ICamp. We then integrated ecDNA/ICamp classification with subclonal architectures for each case (**Fig. 1h**). Many ecDNA-mediated HLAMP regions were clonally amplified, with 11 of 72 (15%) classified as subclonal (amplified in <90% of the cells). In contrast, 143 of 435 (33%) ICamp regions were subclonal, indicating substantial intratumoral heterogeneity. Together, these results establish single-cell CN distributions derived from scWGS as an effective quantitative readout to distinguish ecDNA from ICamp, reflecting their distinct mechanistic modes of inheritance during cell division, and their impact on HLAMP evolution.

### Evolution of ICamps via structural and numeric modulators

Among 435 HLAMP events classified as ICamp, 270 (62%) exhibited multiple CN peaks (**Fig. 2a**), indicating intratumoral heterogeneity (exemplified by case OV-044; **Fig. 2b**). Distinct HLAMP CN peaks generally aligned with phylogenetic clades defined by genome-wide CN profiles. In cases with multiple peaks, individual clones were preferentially concentrated within a single peak rather than distributed across multiple peaks (p<1×10^−16^, Stouffer-combined one-sided permutation test). In aggregate, 69% of subclones were restricted to one peak, consistent with clonal expansions subsequent to subclone-specific ICamp events. Reflecting this heterogeneity, ICamp cases showed significantly greater CN variance than stable genomic regions used as controls (*TP53* and *RB1*; mean standard deviation 3.32 vs. 0.575; p=1.97×10^−72^ by one-sided two-sample *t*-test), although it was significantly less than ecDNA (mean standard deviation: ecDNA 43.1, ICamp 3.32; p=8.68×10^−20^ by one-sided two-sample *t*-test; **Fig. 2c**). To investigate the source of this variability, we classified ICamp events by their generative mechanisms using boundary SV features (**Extended Data Fig. 5a-f**; **Methods**) and asked whether variability differed by initiating mechanism^23,26^. Chromothripsis- and breakage-fusion-bridge (BFB) cycle-mediated ICamps showed greater variance than other groups combined, but the absolute difference was modest (mean standard deviation 3.65 vs. 3.14; one-sided two-sample *t*-test, p=0.0253; **Fig. 2c**), indicating that the initiating mutational processes may have only a small impact on CN variance. To identify processes modulating ICamp CN after their initiation, we compared CN profiles between subclones (**Fig. 2d-e**). We identified pairs of subclones with nearly identical chromosomal or arm-level CN profiles that differed primarily in amplitude, consistent with numeric modulation (4,839 of 8,486 region-subclone pairs, 57%; **Fig. 2f**); these patterns occurred most frequently in the context of whole-genome doubling (WGD; 2,033/4,839, 42%; **Extended Data Fig. 6a**), and the rest from chromosome- or arm-specific aneuploidies (**Extended Data Fig. 6b)**. In other cases, subclones showed dissimilar CN profiles, often accompanied by subclone-specific SVs, indicative of structural modulation (2,939/8,486, 35%; **Fig. 2g**). These involved diverse mechanisms, including subclone-specific fold-back inversion (consistent with additional BFB cycles), chromothripsis, and other complex genomic rearrangements (**Extended Data Fig. 7**).

**Fig. 2.**
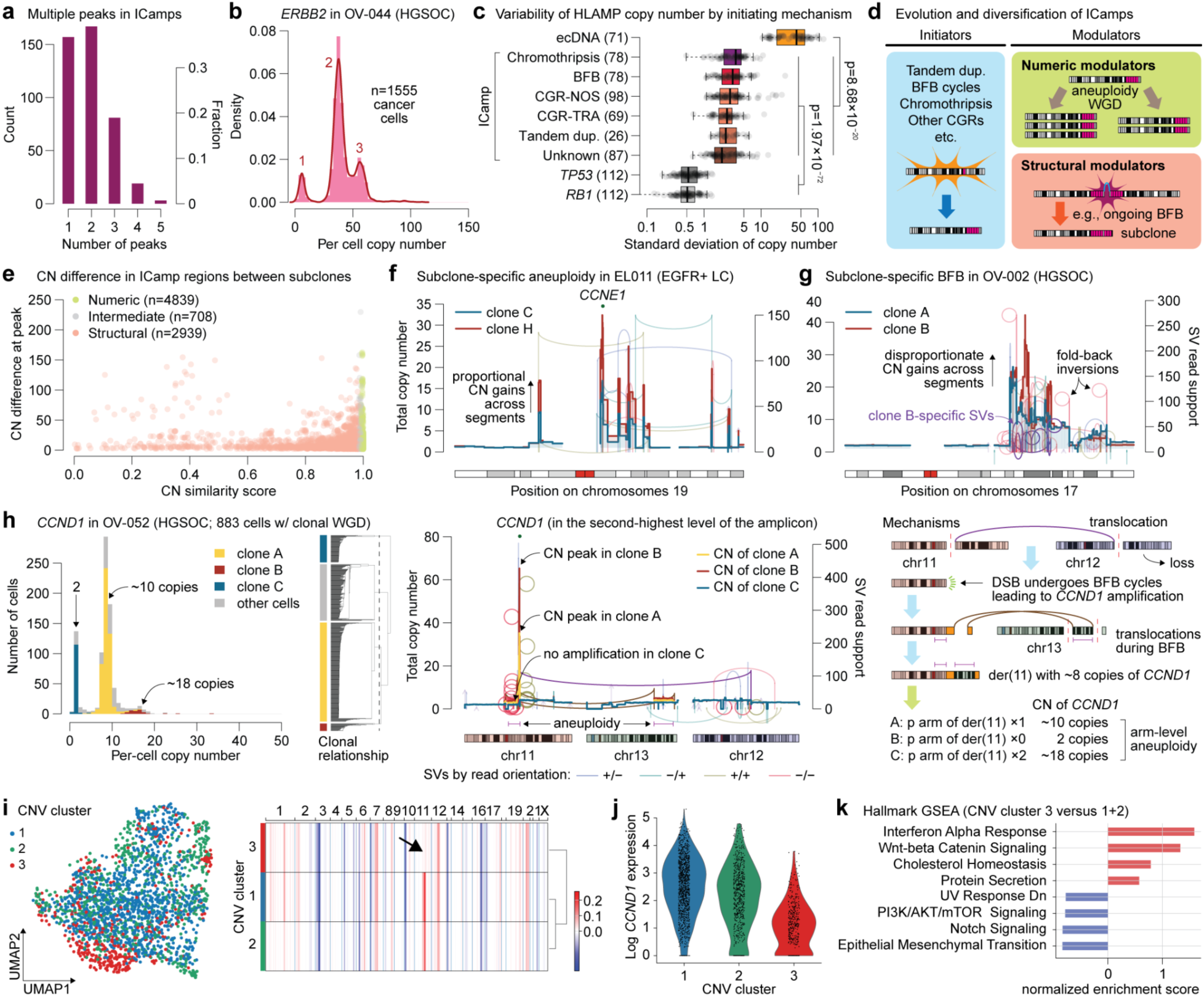
Contrasting mechanisms of ICamp CN heterogeneity by numeric and structural modulators. **a**, Prevalence of multi-peak CN distribution among ICamp cases. **b**, An ICamp case exhibiting multi-peak CN distribution across cells, indicating subclones with distinct *ERBB2* CN. **c**, CN variance of HLAMP events stratified by initiating mechanism (BFB, breakage-fusion bridge; CGR-NOS, complex genomic rearrangement, not otherwise specified; CGR-TRA, complex genomic rearrangement, translocation-rich). **d**, Initiating and modulating mechanisms for ICamps. **e**, Pairwise comparisons of HLAMP-associated arm-level CN profiles between subclones. **f**, A classical example of aneuploidy-mediated doubling of *CCNE1* amplification in EL011. Compared to clone C, all the HLAMP segments exhibit proportional CN increase in clone H. **g**, A structural modulation of 17q amplification in OV-002. Clone B exhibits further amplification of two segments (one amplifying *MIR21*) within the HLAMP segment in clone A. **h**, Multiple CN peaks of *CCND1* amplification in OV-052 matching the subclonal phylogeny (left), clone-specific CNs and SVs indicative of BFB-mediated amplification and subsequent modulation by arm-level aneuploidy (middle), and schematics of the mechanism (right). **i**, UMAP embedding of 2,244 cancer cells from scRNA sequencing of OV-052 (left) and summary CN profiles of inferred clones in the malignant compartment (right). A subclone with no *CCND1* amplification (cluster 3; n=431 cells) is detected. **j**, Normalized expression level of *CCND1* in each cluster. **k**, Gene set enrichment analysis comparing cluster 3 with the other clusters. The top 8 enriched pathways are shown.

As an illustrative example, HGSOC case OV-052 harbored a *CCND1* ICamp with two large and one small CN peaks, indicative of a heterogeneous ICamp. These peaks aligned with three subclones in the CN-based phylogeny (**Fig. 2h, left**). In the major clone A, the *CCND1* amplification, adjacent to an inter-chromosomal rearrangement translocating the distal portion of 11q to 12q, exhibited multiple fold-back inversions at its borders, consistent with BFB cycles further modifying locus subsequent to the translocation. Minor subclone B exhibited arm-level aneuploidy that duplicated the *CCND1* amplicon, reflected by two-fold CN across segments from the centromere through the distal amplicon (**Fig. 2h, middle and right**). Notably, minor clone C lacked evidence of the *CCND1* amplification and showed a subclone-specific loss of the allele with the amplification. Analysis of haplotype-informed allele-specific CN was consistent with arm-level aneuploidy-mediated loss of the *CCND1* ICamp (**Extended Data Fig. 6c**). Moreover, single-cell RNA sequencing from adjacent tumor tissue identified a subclone with negligible cyclin D1 expression and a genome-wide CN profile matching clone C, supporting the phenotypic impact of this aneuploidy-mediated HLAMP loss (**Fig. 2i-j**). Notably, this subclone exhibited a distinct gene expression program, with a high interferon alpha response and upregulation of antigen-presentation machinery (**Fig. 2k**), features proposed to promote an immunosuppressive microenvironment^27^. These results suggest that such bidirectional numeric modulation of ICamp could enhance transcriptional adaptability under selection. We observed similar aneuploidy-mediated elimination of ICamp events in 3 additional cases, often involving G1/S cyclin genes (**Extended Data Fig. 6c**).

We also observed diversification of chromothripsis-mediated HLAMPs through both numeric and structural mechanisms. In one representative case (OV-083), chromosome 8 underwent two rounds of chromothriptic complex rearrangement, generating two subclones that were each further diversified by aneuploidy (**Extended Data Fig. 8a**). In another case (OV-004), chromothripsis of chromosome 8 gave rise to two clades with distinct DNA repair configurations derived from the same intermediate chromosome; as in OV-083, both clades were subsequently diversified by aneuploidy (**Extended Data Fig. 8b**).

Together, our analyses highlight punctuated evolution of ICamp through two principal routes^28^: numeric modulation by aneuploidy and WGD, and structural remodeling via BFB, chromothripsis, and other mutational processes. Notably, these changes occur as both gains and losses and often define sizable subclones, reflective of non-negative fitness and a linkage to tumor evolution.

### ecDNA architecture resolved by cell-to-cell CN variance and genomic rearrangement

We identified 72 HLAMP regions across 11 cases (6 patients and 5 experimental models) with single-cell CN distributions consistent with ecDNA. Notably, genome-wide CN profiles in these 11 cases were relatively homogeneous across cells, with most intercellular differences confined to HLAMP regions. Given the presence of multiple HLAMP segments in each ecDNA-positive case, we first leveraged within-case single-cell CN correlations to infer whether segment pairs were co-located in the same ecDNA species or segregated independently. In GBM0721, two ecDNA-mediated HLAMP segments were identified: one encompassing *MDM4* (1q32.1) and another *EGFR* (7p11.2) (**Fig. 3a, left**). They showed a modest but significant correlation, with considerable variability across the CN range (**Fig. 3a, right**; Pearson correlation=0.25; p=6.5×10^−8^). In contrast, in a surgically resected, heavily pretreated *EGFR*-mutated lung cancer case EL001 where we identified 18 ecDNA-amplified segments, *EGFR* (7q11.2) and *CCND1* (11q13.3) CNs were tightly and linearly correlated despite their different chromosomes of origin (**Fig. 3b**; Pearson correlation=0.98; p=3.4×10^−58^). In GBM0721, the *MDM4* and *EGFR* amplicons likely reside on separate ecDNA species, consistent with independent head-to-tail SVs encircling each segment (**Fig. 3c**). The correlation between *MDM4* and *EGFR* CN may reflect co-selection for a fitness advantage, supported by single-cell RNA-seq from an adjacent tumor region: cells with high *MDM4* and *EGFR* expression showed high proliferative features (e.g., mitotic spindle, Ki-67), whereas cells with low expression of both showed elevated reactive oxygen species signaling (**Extended Data Fig. 9a-c**). By contrast, the linear *EGFR*/*CCND1* correlation in EL001 suggests a chimeric ecDNA species carrying both oncogenes, supported by chromothripsis connecting segments harboring *EGFR* and *CCND1* (**Fig. 3d**). Similarly, *CDK4* and *MDM2* CNs were also tightly correlated (**Extended Data Fig. 10a**) with SV links, implying another ecDNA species co-amplifying these oncogenes. Recognizing that the copy number of segments on the same ecDNA species will co-vary across cells, we computed pairwise correlations among all HLAMP regions in EL001 and performed hierarchical clustering (**Fig. 3e; Methods**). This revealed two largely mutually exclusive correlation clusters, one comprising regions from chromosomes 7 and 11, and another with chromosome 12 regions, consistent with two distinct ecDNA species each harboring two genes.

**Fig. 3.**
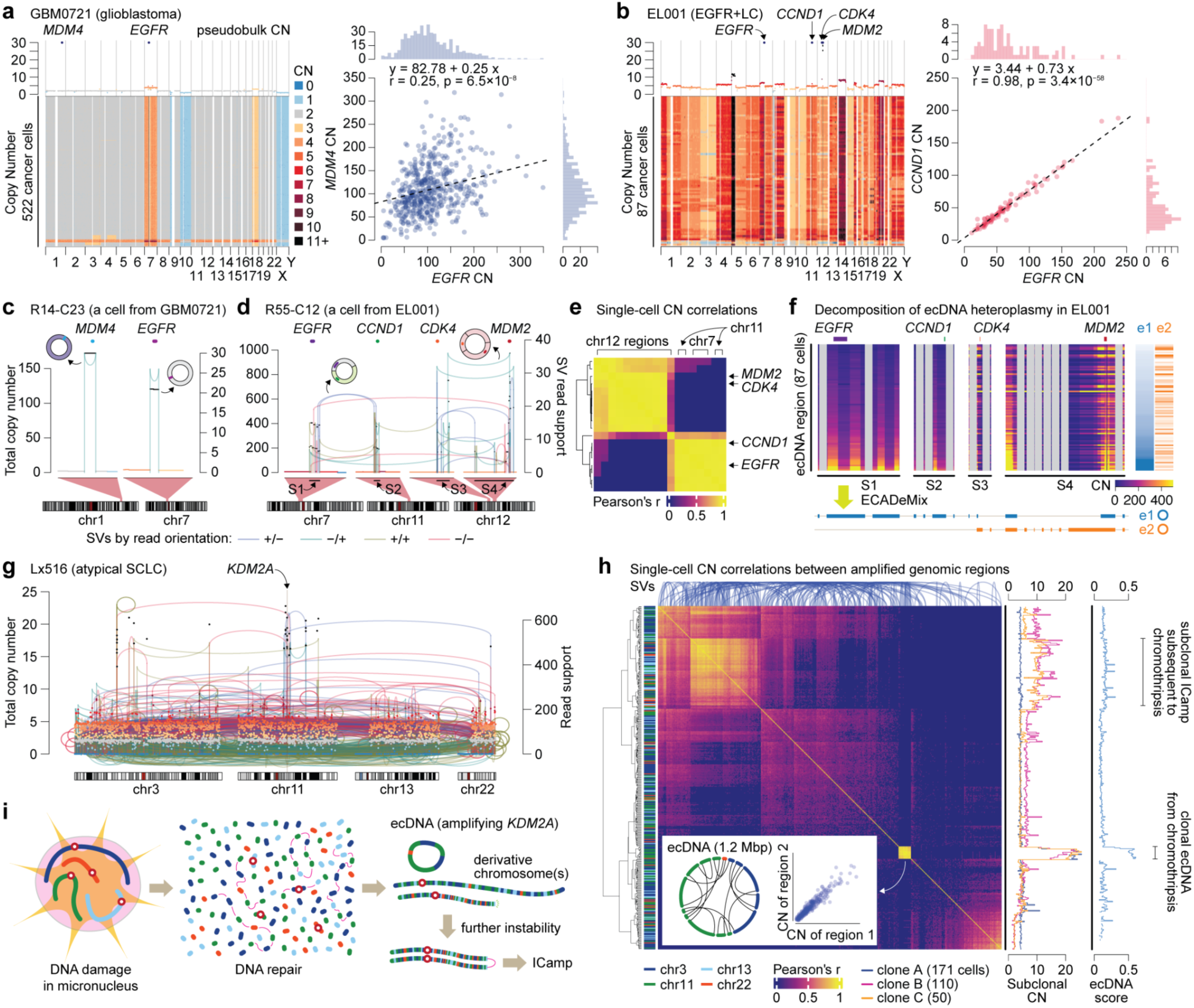
Resolving mechanisms of segregation of co-amplified oncogenes in ecDNA in single cells. **a-b**, Genome-wide CN profiles (left) and CN correlations between key oncogenes (right) for GBM0721 and EL001. **c-d**, Single cell CN/SV profiles indicating two distinct simple ecDNAs in GBM0721 and two distinct hybrid ecDNAs in EL001. **e**, Hierarchical clustering of HLAMP regions in EL001 based on pairwise CN correlations reveals two ecDNA species co-amplifying oncogenes. **f**, Decomposition of ecDNA heteroplasmy in EL001 by ECADeMix. HLAMP segments used (S1-S4) are indicated in **d**. **g**, Pseudobulk CN/SV profile from Lx516 showing multi-chromosomal chromothripsis leading to HLAMP of *KDM2A*. **h**, Clustering based on pairwise CN correlation identifies a hybrid ecDNA and a subclonal ICamp event. Rearrangements between segments are illustrated as arcs on the top. Chromosomal origins in the left heatmap. Subclonal CN and ecDNA scores on the right side. Inset plots show the reconstructed ecDNA structure (left) and a representative CN correlation between two segments that comprise the ecDNA. **i**, Schematic illustration of HLAMP mechanisms in Lx516.

Motivated by these observations, we developed ECADeMix (<u>E</u>xtra-<u>C</u>hromosomal <u>A</u>mplicon <u>De</u>-<u>Mix</u>ing) to systematically identify the structure of ecDNA species and their heteroplasmy across cells. Briefly, analogous to heteroplasmy analyses in mitochondrial DNA in single cells^29^,

ECADeMix jointly infers integer CN profiles for each ecDNA species and their multiplicities per cell by decomposing single-cell CN profiles of ecDNA regions: here to 10-Kbp resolution (**Extended Data Fig. 10-12**; **Methods**). ECADeMix identified two ecDNA species in each of GBM0721 and EL001, confirming our previous conclusions and additionally revealing that in EL001 the *MDM2* locus was also present on the *EGFR*/*CCND1* ecDNA (**Fig. 3f**), a feature that could not be resolved from correlation analyses. Furthermore, ECADeMix identified ecDNA species co-amplifying multiple oncogenes in two other cases (*MYC* and *CDX2* in COLO320-DM and *NKX2-1* and *FOXA1* in EL003) as well as those in which multiple distinct clone-specific species separately amplified the same oncogene (e.g., GBM0510, discussed below). Using ECADeMix, we identified a total of 23 ecDNA species across 11 cases, including 6 cases with multiple species (2-11 each). This included surgically resected brain metastases of atypical SCLC characterized by an exceptional multi-chromosomal chromothripsis involving four chromosomes, resulting in ecDNA-mediated HLAMP (**Fig. 3g**). Given the extreme complexity of this case, we questioned how many ecDNA species had formed through this process. Notably, ECADeMix identified only a single ecDNA species conjoining 13 regions originating from 3 chromosomes (**Extended Data Fig. 10d**). Accordingly, pairwise CN correlations among all 391 amplified regions in this case (152,881 pairs) revealed a distinct cluster of near-perfect correlations encompassing all 13 regions that formed a full circle in structural reconstruction (**Fig. 3h)**. Additional regions showed strong but more modest correlations (left upper part of the heatmap); their CN profiles and ecDNA scores are consistent with subclonal ICamp events arising from chromothriptic derivative chromosome(s), likely reflecting ongoing instability (**Fig. 3i**). Together, these analyses illustrate that cell-to-cell CN variance and pairwise segment correlations enable deconvolution of complex ecDNA architectures. Further, ECADeMix analysis of scWGS unveils new evolutionary models for ecDNA through robust estimates of both presence and prevalence of ecDNA species in each cell, enabling an algorithmic approach to dissect complex ecDNA-mediated cancer evolution.

### ecDNA evolution at single-nucleotide resolution

We next integrated SV features and amplicon species information at the single-cell level to elucidate how ecDNA drives cancer evolution. The high coverage of ecDNA regions enabled single cell-level, nucleotide-level analysis of SVs (median per-cell coverage 11.09X, range 0.9522X−48.11X). In 5 of 11 ecDNA-positive cases (Lx33, COLO320-DM, GBM39-DM, NCI-H69, NCI-H524, 2765_2), we detected a dominant ecDNA species accompanied by subpopulation of cells sharing lower-frequency variant forms, consistent with branched (divergent) evolution that followed initial clonal expansion by the ancestral ecDNA (**Extended Data Fig. 11a-c**). For example, the NCI-H69 SCLC cell line harbored a chromothripsis-derived ecDNA amplifying *MYCN* (**Fig. 4a, top left**) with more than half of cells carrying a predominant parental ecDNA species (e0) containing all chromothriptic HLAMP segments. Using ECADeMix, we identified three variant ecDNA forms (e1-e3) exhibiting internal CN losses in distinct regions, consistent with derivative forms of e0 (**Fig. 4a, bottom left**). These variant ecDNAs coexisted with e0 in some cells, providing further support for their derivation from the parental molecule. Single-cell SV analysis identified specific rearrangements that account for the variant-specific internal deletions (**Fig. 4a, right**). We considered whether these deletions reflect active exclusion of deleterious genetic elements; in NCI-H69 and NCI-H524, the two samples where the variant ecDNAs are most prevalent, this seems unlikely as the deletions predominantly affect gene deserts or already-truncated genes (**Extended Data Fig. 12a-b**). Instead, the data are more consistent with ongoing ecDNA remodeling shaped by stochastic DNA damage and repair^30^, with potential purifying selection preserving segments harboring essential oncogenic elements, such as *MYCN* (**Fig. 4b**). In addition, we identified cells exhibiting unique CN patterns in ecDNA regions, suggesting ongoing branched evolution (**Extended Data Fig. 12c**).

**Fig. 4.**
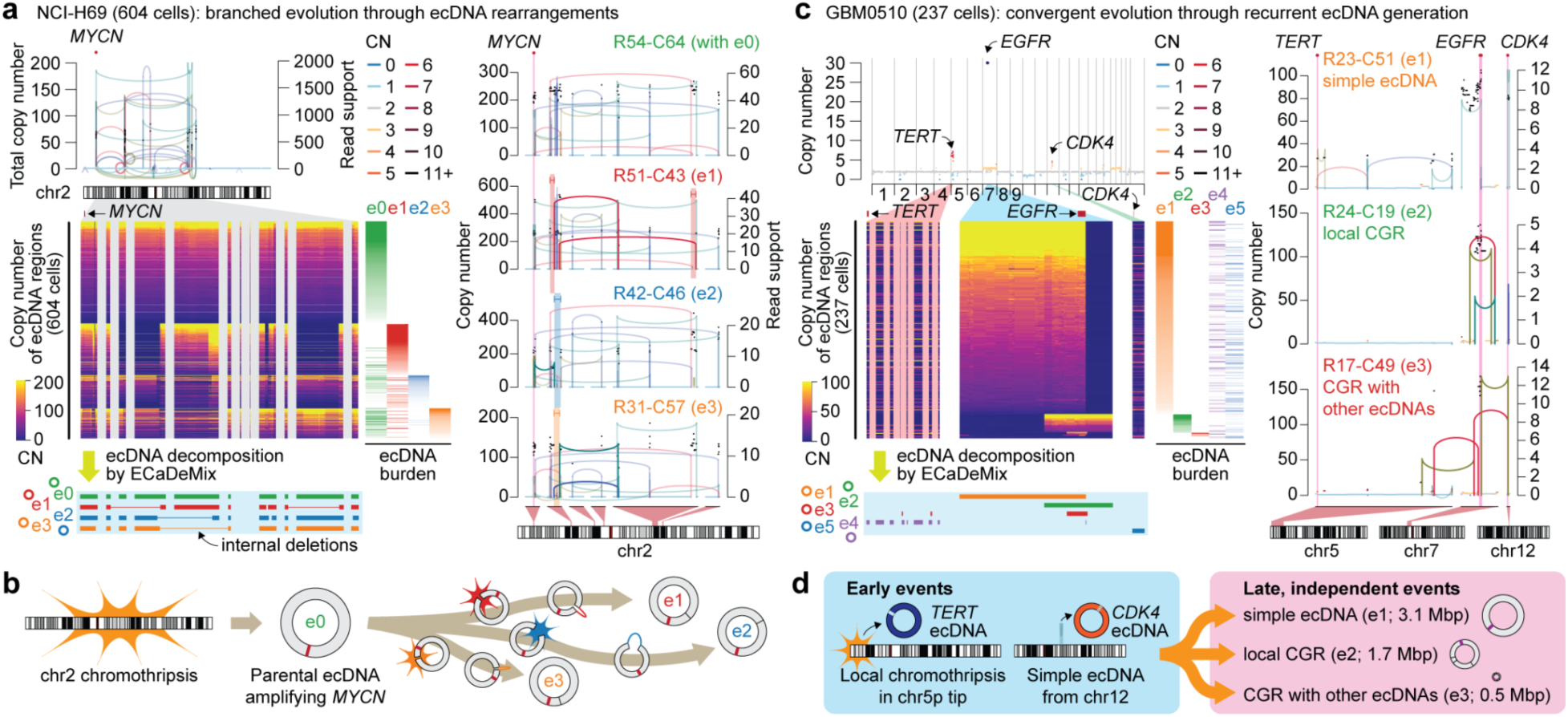
Modes of ongoing ecDNA evolution. **a**, Chromothripsis-derived ecDNA amplifying *MYCN* in the NCI-H69 SCLC cell line (top left); single-cell CN profiles of the amplicon and ecDNA heteroplasmy delineated by ECADeMix (bottom left). CN/SV profiles from 4 cancer cells with distinct ecDNA subspecies (right). e0 is the likely parental ecDNA; e1-e3 are daughter ecDNAs derived from e0 via internal rearrangements. **b**, Schematic illustration of branched evolution via ecDNA rearrangement. **c-d**, Pseudobulk genome-wide CN profile of glioblastoma GBM0510, exhibiting multiple ecDNA-mediated HLAMPs (**c,** top left); single-cell CN profiles of the amplicons and decomposed ecDNA species (**c,** bottom left). *EGFR* is amplified by three distinct ecDNAs (e1, e2, and e3) in a mutually exclusive manner across cells, forming distinct subclones (single-cell examples in the right panel of **c**). ecDNAs amplifying *TERT* and *CDK4* are monotypic, likely reflecting temporal order among the events.

In contrast, GBM0510 provides evidence of convergent evolution^31^ through ecDNA. In this tumor, ECADeMix revealed that three oncogenes—*TERT* (5p15.33), *EGFR* (7p11.2), and *CDK4* (12q14.1)—were each amplified on separate ecDNA species (**Fig. 4c, top left** and **Extended Data Fig. 13a**). *TERT* and *CDK4* ecDNAs were subclonal, present in 47% and 46% of cells, respectively, and were structurally uniform within positive cells: *TERT* ecDNA arose from a local chromothripsis event in the subtelomeric region 5p region, and *CDK4* ecDNA formed by a simple head-to-tail rearrangement (**Fig. 4c, bottom left**). In contrast, *EGFR* amplification was clonal but partitioned into three mutually exclusive ecDNA forms (e1-e3) across cells. The largest and most prevalent (e1) was formed by a simple head-to-tail rearrangement, whereas e2 and e3 were generated by distinct, complex rearrangements (**Fig. 4c, right**). The amplicon intervals for e1-e3 overlap only at the *EGFR* locus, and no SVs are shared among them, supporting independent origins. Single-nucleotide variant (SNV) profiles (**Extended Data Fig. 13**) indicate that the subclones defined by the three *EGFR* ecDNA forms share most SNVs and harbor few private SNVs, consistent with relatively late emergence of these ecDNA forms^32^. Taken together, these features support convergent evolution through recurrent, independent acquisition of *EGFR*-amplifying ecDNAs. Compared with the multiple *EGFR* ecDNA species, *TERT* and *CDK4* ecDNAs were structurally uniform, consistent with earlier acquisition, whereas their subclonality may reflect stochastic loss from unequal ecDNA segregation (**Fig. 4d**). Similarly, in GBM0721 we identified three mutually exclusive ecDNA species amplifying *EGFR*, one of which predominated (**Extended Data Fig. 13b**). In contrast, we found only a single ecDNA species for *MDM4*, again supporting a late acquisition of ecDNA-mediated *EGFR* amplification in GBM. Thus, these findings are consistent with ecDNAs in human cancers mediating both divergent and convergent evolution, resulting in dynamic cell-to-cell diversification over time.

### ecDNA status by scWGS relative to bulk genome-graph predictions

We next asked how single-cell analyses of ecDNA species and their evolutionary trajectories compare to bulk genome-graph based predictions. For this comparison, we applied Amplicon Architect (AA), a widely used method for identifying ecDNA from bulk sequencing data^33^, to pseudobulk data generated from our single-cell WGS (**Fig. 5a; Methods**). We then matched HLAMP regions to AA-defined amplicons and compared their classifications. Importantly, pseudobulk data derived from our scWGS are effectively analogous to bulk WGS but without information on CN distributions over cells, enabling a head-to-head comparison of the two approaches. At the case level, 9 out of 11 ecDNA-positive cases by single-cell CN distribution were also called ecDNA-positive by AA (81% concordance). However, among 82 cases without a CN distribution consistent with ecDNA, AA called 30 (37%) as ecDNA-positive. At the HLAMP region level, stratified by cancer type (**Fig. 5b**), discrepancies were mainly in HGSOC and hormone receptor-negative (HR−) breast cancers: these regions exhibited one or more narrow, high CN peaks in single cells, indicating ICamp (median ecDNA score=0.129, range=0.040−0.380). In contrast, ecDNA-positive cases by AA in *EGFR*-mutated NSCLC, SCLC, and GBM mostly showed CN distributions consistent with ecDNAs, suggesting tissue-type specificity of ecDNA prevalence. Review of AA genome graphs suggested several potential sources of discrepancies (**Extended Data Fig. 14a-c**). In ecDNA-positive cases by scWGS, the CN transitions between unamplified and amplified segments were typically abrupt, consistent with circularization followed by amplification, and exhibited high junctional CN in pseudobulk profiles (**Fig. 5c** and **Extended Data Fig. 14a**)^34^, with notable exceptions in cases with subclonal ecDNAs. However, in many HLAMP cases in HGSOC and HR− breast cancers, pseudobulk CN profiles showed stepwise increases in CN from unamplified to amplified regions across extended intervals, despite SVs arranged in cyclic paths (**Extended Data Fig. 14b**). This pattern is more consistent with progressive ICamp via multiple rearrangements, than with ecDNA^35^.

**Fig. 5.**
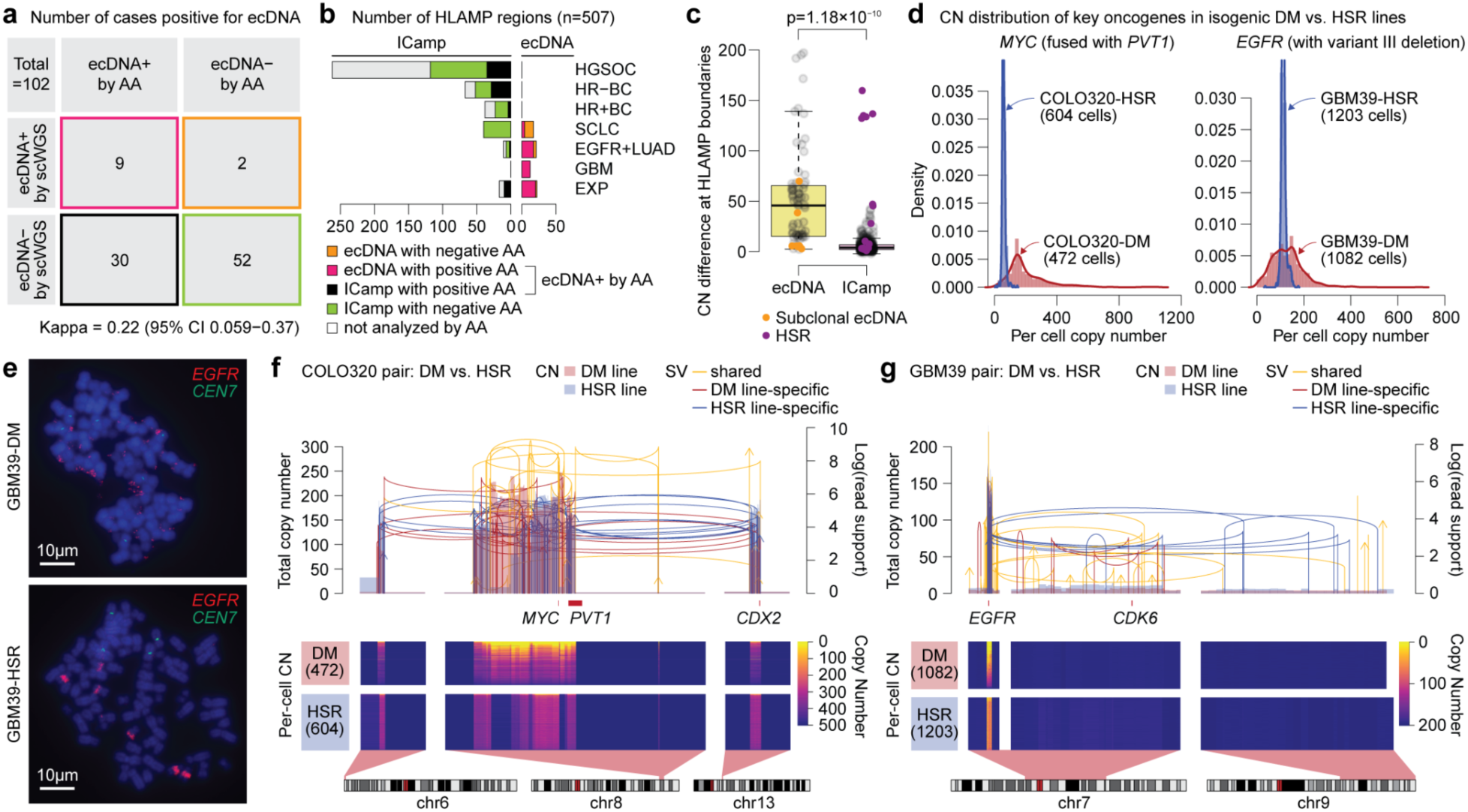
scWGS distinguishes current ecDNA from historical bulk-level footprints. **a**, Case-level comparison of ecDNA predictions from bulk genome-graph methods (Amplicon Architect/Classifier) versus scWGS. **b**, HLAMP region-level comparison divided by cancer type. **c**, Junctional CN difference profiles in ecDNA and ICamp regions. Statistical significance was determined by Student’s *t* test. **d**, CN distributions in isogenic paired cell lines: ecDNA-mediated double minutes (COLO320-DM and GBM39-DM) versus homogeneously staining regions (COLO320-HSR and GBM39-HSR). **e**, Cytogenetic features of GBM39 isogenic pair exhibiting numerous variable sized, dispersed signals consistent with *EGFR*-bearing ecDNA (top) and intrachromosomal clustered amplification manifesting as homogeneously staining region (HSR). **f-g**, Pseudobulk CN/SV profiles (top) and single-cell CN profiles (bottom) for DM versus HSR lines reveal genomic rearrangements that occurred after the most recent common ancestor and prior to HSR formation in COLO320 and GBM39 isogenic pairs.

Notably, three HLAMP cases showed circular paths in AA genome graphs and abrupt CN transitions in pseudobulk profiles, a combination typical for ecDNA, yet their single-cell CN distributions were instead consistent with ICamp (**Fig. 5c**). Two of these were isogenic cell line models exhibiting homogeneously staining regions (HSRs) in DNA FISH (COLO320-HSR and GBM39-HSR; **Fig. 5d-e**)^36,37^. Similar to their ecDNA-positive counterparts (COLO320-DM and GBM39-DM, respectively), these HSR cell lines retain the highly amplified SV breakpoint footprint of the initiating ecDNA (**Fig. 5f** and **5g**), indicating that the HSR was formed by chromosomal integration of ecDNAs^38^. The third case was NCI-H82, a SCLC cell line, which was labeled as ecDNA-positive in a previous study using AA^39^. The bulk genome graph showed a cyclic path amplifying *MYC* with abrupt CN transitions, but the single-cell CN distribution was sharply peaked, consistent with ICamp (**Extended Data Fig. 14c**). DNA FISH confirmed *MYC* HSR formation in this line without evidence of ecDNA (**Extended Data Fig. 4b**).

Last, we analyzed genomic rearrangements associated with chromosomal integration of ecDNA in isogenic pairs. In COLO320-HSR, the core amplicon contained several internal deletions indicated by large, abrupt CN transitions, consistent with branched evolution from the parental ecDNA prior to integration (**Fig. 5f**). In GBM39-HSR, HSR line-specific SVs included a chromothripsis event linking the *EGFR* amplicon to the subtelomeric 9q region (**Fig. 5g**). These HSR line-specific SVs inform mutational processes between the shared ecDNA ancestor and HSR formation but do not, by themselves, identify the integrated chromosomes. Orthogonal mapping from previous studies places the HSR on chrX for COLO320-HSR^40^. Together, these analyses show that single-cell CN distributions can distinguish actively maintained ecDNAs from historical ecDNA footprints re-integrated into chromosomes, substantially revise bulk genome-graph predictions, and place HLAMP events within their evolutionary context.

## Discussion

Leveraging scWGS, we quantified cell-to-cell variability in HLAMP across four major cancer types and experimental models, and linked this variability to specific generative and modulatory mechanisms. In contrast to bulk genome-graph predictions, single-cell CN distributions of HLAMP events distinguish episomal ecDNA from ICamp, including HSRs derived from chromosomal re-integration of ecDNAs. Although cyclic genome graphs in bulk WGS can nominate candidate ecDNAs, many candidates, particularly in HGSOC and HR− breast cancers, show narrow, sharply peaked CN distributions consistent with ICamp. In these tumor types, HLAMP heterogeneity appears largely shaped by ICamp evolution through numeric modulation and structural remodeling, which can generate substantial intratumoral heterogeneity in a punctuated manner. Notably, subclones that lose oncogene amplification through aneuploidy warrant further clinical investigation, as selection for oncogene-low subclones could promote early resistance to oncogene-directed targeted therapies^41^.

In contrast to enrichment-based single-cell methods that selectively profile ecDNA^42–44^, scWGS enables genome-wide characterization of HLAMP evolution in single cancer cells, including both ecDNAs and ICamps. Compared with other genome-wide single-cell profiling methods such as single-cell ATAC^45^ and RNA sequencing^46^, which are being explored for ecDNA detection, DLP+-based scWGS or related technologies^47,48^ provide unbiased CN profiles and SV information with high CN fidelity and nucleotide-level coverage at HLAMP loci in single cells, both of which are crucial for defining the mechanisms and evolution of HLAMP events. Heuristically, in the analyses presented here, we required at least 30 high-quality single-cell CN profiles for ecDNA detection. However, we suggest further algorithmic improvements and context-specific events with ultra-high copies could be resolved to fewer than 30 cells.

Our results suggest that ecDNA is prevalent in select cancer types, in contrast to recent studies based on bulk genome-graph predictions^49,50^. In cancer types where ecDNA has been extensively described, namely GBM and SCLC, our scWGS-based analysis confirms a high prevalence of ecDNA. We also identified 3 ecDNA species in 2 of 5 *EGFR*-mutated NSCLCs, amplifying key oncogenes including *EGFR*, *MDM2*, and *NKX2-1*. The contribution of ecDNAs to the biology of *EGFR*-mutated lung cancer and to responses to targeted therapies will be of clinical interest. By contrast, we did not observe any HLAMP event with a CN distribution consistent with ecDNA among 72 patients with HGSOC or HR− breast cancer. Although cyclic paths were identified around HLAMP regions in many cases, their single-cell CN distributions and SV features were instead consistent with ICamps arising through multi-step rearrangement processes. In some cases, rare cells displayed CN outliers superimposed on a global CN distribution consistent with ICamp. These outlier cells may contain ecDNA, but their frequency was exceedingly low. The striking paucity of ecDNA in HGSOC and HR− breast cancers suggests that these tissue contexts could be less permissive to ecDNA formation or maintenance. In addition, previous studies indicated that anticancer therapies, such as chemotherapy or radiotherapy, can induce ecDNA formation^51,52^, potentially reshaping ecDNA prevalence in patients with advanced cancer after treatment. In line with this, a clinical study in patients with platinum-resistant HGSOC reported double minutes in cancer cells from patients’ ascites^53^. The interplay between tissue context and therapy-induced genome instability in shaping the HLAMP landscape warrants further investigation.

Motivated by longstanding efforts to develop ecDNA-directed therapeutics^13–15,54,55^ and growing recognition of the profound impact of ecDNA on cancer evolution, there is increasing interest in detecting ecDNAs in patient biospecimens. Conventional cytogenetic techniques like FISH require metaphase cells to accurately detect ecDNAs^56^, but these are not often available for clinical tumor samples. Instead, high-throughput sequencing of clinical tumors has become an integral part of oncology care, and the use of such assays to detect ecDNA has been proposed^8^. Our findings highlight intrinsic challenges to this approach. A key genomic hallmark of prevalent ecDNA is pronounced cell-to-cell CN variability, which cannot be resolved from single-sample bulk sequencing. Genome graph-based approaches instead depend on sensitive detection of SVs from WGS and cannot reliably distinguish extant ecDNA from its vestigial footprints after chromosomal integration. Given the dynamic nature of HLAMP evolution, including the widespread punctuated modulation of ICamp events observed in our study, scalable scWGS, or targeted single cell approaches whereby cellular distributions can be read out on specific loci, may offer a more promising path for clinical ecDNA detection and monitoring in the future.

In conclusion, our analysis of >86,000 single cell cancer genomes defines interpretable CN distributions of HLAMP events consistent with distinct ICamp and ecDNA generative processes and elucidates how these mechanisms diversify cancer cell populations. These dynamics extend beyond ecDNA and are pervasive in tumors with ICamp, underscoring ongoing genome remodeling. Single-cell genomic profiling of longitudinal clinical specimens, integrated with orthogonal readouts of gene expression^48^, would enable direct measurement of how therapy reshapes HLAMP evolution. The derivative single-cell readouts of architecture, abundance and clonal selection in cancer cells will therefore help to identify actionable dependencies and inform therapeutic strategies through an evolutionary lens.

## Acknowledgments

This work was supported by the Halvorsen Center for Computational Oncology and Cycle for Survival supporting Memorial Sloan Kettering Cancer Center. S.P.S. holds the Nicholls Biondi Chair in Computational Oncology and is a Susan G. Komen Scholar. Additional support from the Department of Defense Congressionally Directed Medical Research Program award [W81XWH-20-1-0565], an Ovarian Cancer Research Alliance (OCRA) Collaborative Research Development Grant [648007], an NIH R01 CA281928-01, and by the Seidenberg Family Foundation funds to SPS. Additional support was provided through the MSK Core Grant (P30 CA008748), the National Cancer Institute of the National Institute of Health (K08CA301011 to JJL; K99CA277562 to SS; R01CA282913 to AV), the Burroughs Wellcome Fund Career Award for Medical Scientists (to JJL), the Cancer Grand Challenge Initiative (to AV), the American Cancer Society Discovery Boost Grant (to AV), the Mark Foundation for Cancer Research ASPIRE Grant (to AV). We thank Gryte Satas, Ignacio Vazquez-Garcia, Samuel Freeman, Matthew Zatzman, Alessandro Grande, Giovanni Santacatterina, and the rest of the Shah, McPherson, and Lee labs for discussions and technical advice.

## Author Contributions

JJL and SPS conceived the project and supervised study design. JJL and SS designed experiments and performed primary computational analyses, with conceptual input from MJW and AM. SS led development of the amplification classifier eicicle. MAM led development of the ecDNA deconvolution algorithm ECADeMix. SS performed single-cell RNA sequencing analyses. MJW conducted allele-specific copy-number analyses. MY, DHA, EGS, and AV performed cytogenetic experiments. MY and JL conducted cell culture experiments. PM, ER, AQV, and CMR generated data from patient-derived xenograft models. PR, SA, N. Rekhtman, VT, MMA, HAY, KKHY, and CMR generated data from clinical tumor samples. JJL, SS, MAM, MJW, KT, CT, CZJP, SM, and AM performed secondary computational analyses. SC, EH, and APS established and maintained computational analysis pipelines. MW and NM supervised single-cell whole-genome sequencing procedures. JJL, SS, MAM, N. Rusk, and SPS wrote the original draft, and all authors revised and approved the final manuscript.

## Competing Interests

A patent application related to the determination of ecDNA versus ICamp based on single-cell DNA CN information has been filed by Memorial Sloan Kettering Cancer Center. AQV is a current employee of AstraZeneca. PR reports research funding from Grail, Novartis, AstraZeneca, EpicSciences, Invitae/ArcherDx, Biothernostics, Tempus, Neo- genomics, Biovica, Guardant, Personalis, Myriad and consulting or advisory role for Novartis, AstraZeneca, Pfizer, Lilly/Loxo, Prelude Therapeutics, Epic Sciences, Daiichi-Sankyo, Foundation Medicine, Inivata, Natera, Tempus, SAGA Diagnostics, Paige.ai, Guardant, and Myriad, outside the scope of this work. SA is cofounder and shareholder of Genome Therapeutics and reports advisory roles for Chordia Therapeutics and Sangamo Therapeutics, outside the scope of this work. MMA reports research funding from Amgen, AstraZeneca, Bristol Myers Squibb, Genentech and Lilly, consulting or advisory roles for Affini-T, AstraZeneca, Blueprint Medicines Corporation, D3Bio, EMD Serono, Gritstone, Iovance, Merck, Merus, Mirati, Novartis, Pfizer and Synthekine, and serves on the Data Safety Monitoring Boards for Apollomics and Bristol Myers Squibb, outside the scope of this work. HAY reports consulting or advisory role for Takeda, Taiho, Black Diamond, BMS, AbbVie, Amgen, AstraZeneca, Daiichi Sankyo, Ipsen, and Pfizer and serves on the Data and Safety Monitoring Board for Janssen and Mythic Therapeutics, outside the scope of this work. CMR reports consulting or advisory role for Genentech/Roche, AstraZeneca, Amgen, Jazz Pharmaceuticals, Earli, AbbVie, Daiichi Sankyo/UCB Japan, Merck, Auron Therapeutics, DISCO and research funding from Merck, Genentech/Roche, Daiichi Sankyo, outside the scope of this work. SPS reports research funding from AstraZeneca and Bristol Myers Squibb, outside the scope of this work.

## Data Availability

Newly generated raw scWGS data will be available from the NCBI dbGaP prior to publication. Previously published raw scWGS datasets are available through European Genome-Phenome Archive (references 17 and 19: EGAS00001006343 and EGAS00001003190) and dbGaP under the National Institute of Health (references 20 and 21: phs002857.v3.p1).

## Code Availability

Source code for the computational models and scripts used for analysis in this manuscript can be found at https://github.com/shahcompbio/hlamp.

## Methods

### Study Cohort

Our study cohort comprises both previously published single-cell whole-genome sequencing (scWGS) datasets from our group and collaborators and newly generated scWGS data produced for this study. All scWGS data were generated using the direct library preparation plus (DLP+) platform.

For published data, we assembled a meta-analysis cohort from 4 previous studies^17, 19–21^, including surgically resected high-grade serous ovarian cancer (HGSOC) samples from 46 patients, patient-derived xenograft (PDX) models of HGSOC from 16 patients, and PDX models of hormone receptor-negative breast cancers from 10 patients (9 were also negative for *HER2* amplification and 1 *HER2*-amplified). PDX-based scWGS data was generated at the British Columbia Cancer Research Centre, whereas scWGS data from clinical HGSOC tumor samples were generated at Memorial Sloan Kettering Cancer Center (MSKCC).

We additionally generated new DLP+-based scWGS data for this study from 3 cancer subtypes—small-cell lung cancer (SCLC), *EGFR*-mutated non-small cell lung cancer (NSCLC), and glioblastoma (GBM)—as well as several experimental models. For SCLC, we profiled PDX models established from 5 patients and rapid autopsy-based tumors from 3 participans. For *EGFR*-mutated NSCLC, we used frozen surgical tumor resections from 5 patients. For GBM, we used freshly dissociated single-cell suspensions derived from surgical resection from 4 patients. These samples were processed at the Integrated Genomics Operation at Memorial Sloan Kettering Cancer Center.

All patients provided informed consent under the Institutional Review Board (IRB)-approved clinical study protocols. The study was conducted in accordance with the Declaration of Helsinki and Good Clinical Practice (GCP) guidelines.

### DLP+ Analyses

#### ScWGS Data Alignment and Quality Control

We processed scWGS data as described previously using the publicly available Mondrian software suite (https://github.com/mondrian-scwgs/mondrian), executed within the Isabl framework^57^. After trimming adaptor sequences, reads from each cell were aligned to the human reference genome GRCh37 using bwa-mem v0.7.17. PCR duplicates were identified and marked using Picard v2.27.4. Alignment and quality metrics were collected from each BAM file using Picard (InsertWgsMetrics and WgsMetrics) and samtools flagstat. BAM files from each cell were then merged to generate a pseudobulk BAM file.

For first-pass inference of single-cell copy-number profiles, reads were counted in non-overlapping 500-Kbp bins genome-wide for each cell. GC-content bias was adjusted using modal regression normalization. Copy-number states were then inferred using HMMcopy^58^ by selecting the best-fit model.

Finally, we integrated alignment metrics and copy-number profiles through the quality-control pipeline to estimate quality of each cell and to flag S-phase cells and doublets. Based on this result, we filtered non-cancer cells (either control cells or those exhibiting euploid copy-number profiles), low-quality cells (quality score less than 0.75), cells with low read count (total mapped reads ≤ 250,000), S-phase replicating cells, and doublets, resulting in 86,239 high-quality single-cell cancer genomes.

#### Somatic Structural Variant Analysis

We detected somatic structural variants (SVs) using GRIDSS v2.13.2^59^ with default parameters and hg19 blacklisted regions from the ENCODE consortium (ENCFF001TD0; https://www.encodeproject.org/files/ENCFF001TDO/). For the 82 cases where bulk whole-genome sequencing (WGS) of normal tissue is available, we ran GRIDSS in multi-sample mode, treating all DLP+ libraries as tumor samples and the matched normal bulk WGS as the normal. For the remaining 18 cases without matched normal WGS (**Table S1**), we ran GRIDSS in tumor-only mode. VCF output files were post-processed to collapse multi-sample SV calls into a single composite column. The resulting composite VCF was then subject to GRIPSS v.2.3 (https://github.com/hartwigmedical/hmftools/tree/purple-v3.7.2/gripss) for somatic filtering. GRIPSS was run with default settings, using the panel of normals, fusion hotspot, and repeat information recommended by the Hartwig Medical Foundation tool suite. The resultant VCF file was converted into BEDPE format for further downstream analysis (associated scripts are available at: https://github.com/shahcompbio/hlamp/tree/main/src/ <u>structural_variant_calling</u>).

We next genotyped the SVs detected at pseudobulk level in each single cell using a modified version of SVtyper (https://github.com/marcjwilliams1/svtyper)^60^, as described in our recent study^21^. In brief, we modified SVtyper to compute the number of read support in each single cell. It provides per-cell read support for the reference and variant sequences, including split reads that directly capture the breakpoint and discordant read pairs with larger-than-expected insert sizes with proper orientation (+/−) or those in variant orientation (−/+, −/−, +/+) or mapping to different chromosomes. We added features to output cell IDs as concatenated string in the VCF and generate matrices linking cell IDs and SV IDs for downstream analysis.

#### SV Breakpoint-Concordant Single-Cell Copy-Number Analysis

To infer the generative mechanisms of high-level copy-number amplification (HLAMP) events from associated SV features, we generated case-specific genomic segments by integrating SV breakpoints with a genome-wide tiling of non-overlapping 2 Mbp bins. Using pseudobulk BAM file, we then computed per-cell read counts for each segment in all high-quality cancer cells (https://github.com/shahcompbio/hlamp/tree/main/src/sv_aware_seg_cn/02_ <u>sv-seg-cn/</u>). Then, we computed the average reads per copy per base in the HMMCopy 500kb copy number profiles and used this constant for each cell to scale the segment read counts to approximate segment copy numbers (https://github.com/shahcompbio/hlamp/tree/main/src/sv_aware_seg_cn/03_calibrate_cn_gc_cell_inclusion.py). This scaling ensures that despite varying segment sizes, the approximate copy numbers would be comparable between segments. Next, we applied GC correction via modal quantile regression^16^. Finally, we scaled these GC-corrected values to match the cell ploidy estimated using HMMCopy (assuming that any HLAMPs whose copy number is now captured more accurately would not meaningfully contribute to the cell ploidy due to their small size).

#### Clustering Cancer Cells to Identify Subclones

High-quality cancer cells were clustered based on their genome-wide copy number profiles. Briefly, we first restricted the bin set to informative loci by removing bins with zero normalized read coverage, excluding short segments (<1000 bp), and removing bins overlapping centromeric regions. We then constructed a cell by bin matrix from the GC-corrected copy number layer, set any non-finite values to zero, and standardized features per bin using z-scoring with clipping of extreme values (maximum absolute value 10). Dimensionality reduction was performed by principal component analysis (PCA; up to 50 components), followed by construction of a k-nearest neighbor graph (k=15) in PCA space and community detection using the Leiden algorithm at resolution 0.7 (unless otherwise specified). These subclones were labeled by the Leiden partition. This procedure was followed by generating case-specific diagrams (https://github.com/shahcompbio/hlamp/tree/main/src/sv_aware_seg_cn/04_dlp.composite.plot. <u>v4.R</u>) including pseudobulk copy number plot, per-cell copy number heatmap, and subclone-specific copy-number plots for manual review.

#### Identifying HLAMP regions

Using signals package^17^, we first estimated pseudobulk and subclonal copy-numbers based on the SV breakpoint-concordant single-cell copy-number profiles and subclones defined above. Then we established baseline copy number for each chromosome arm at pseudobulk level, by selecting the copy-number state with the greatest combined segmental length, with the lowest possible baseline copy number value being 2, excluding acrocentric arms (13p, 14p, 15p, 21p, and 22p). We then marked the segments of which copy number value is more than 3 times of the arm-level copy-number baseline in at least one subclone and combined contiguous segments to define amplified regions. For those regions that were bordered by fold-back inversions (SVs of which orientation is −/− or +/+ and the distance between two breakpoints were 5000 bp or less), we calibrated the borders by including both inversion breakpoints (https://github.com/shahcompbio/hlamp/tree/main/src/sv_aware_seg_cn/05_dlp.oncogene.cn.v4.R and https://github.com/shahcompbio/hlamp/tree/main/src/sv_aware_seg_cn/06_dlp.region.cn.v4.R). Applying this process on 102 cases identified 2,925 amplified regions. Among these, 507 segments of which weighted mean copy number is 6 or greater, we labeled them as HLAMP and used as main data for downstream analyses.

#### Mechanistic classification of HLAMP regions based on SV features

SVs at boundaries between amplified and un-amplified regions can provide insights into the generative mechanisms of an amplicon^23,61^. Our SV breakpoint-aware single-cell copy-number profiles enabled classification of HLAMP events into mechanistic classes based on SV patterns at HLAMP boundaries.

We first clustered the SV breakpoints using clusterSV (https://github.com/cancerit/ClusterSV), which accounts for breakpoint proximity, background SV rates, and SV type-specific size distributions, as described previously^26^. We defined chromothripsis as an SV cluster meeting three criteria: i) ≥20 SV breakpoints on a single chromosome; ii) SVs represented in all 4 read orientations, and iii) unbiased representation across the 4 orientation categories, assessed by permutation testing. For iii), we tested whether the observed counts of SV breakpoints across 4 orientations differed from expectation under equal probability using a chi-square goodness-of-fit test (implemented via the permutation_test function in https://github.com/ <u>shahcompbio/hlamp/tree/main/src/sv_aware_seg_cn/07_ amp.mechanism.R</u>).

HLAMP events whose boundary SVs were part of chromothripsis cluster were classified as **chromothripsis**. HLAMP events bordered by fold-back inversions (FBIs) were classified as **FBI**. If both boundaries were connected by a single head-to-tail SV, the event was classified as **tandem duplication**. If neither boundary had a detected SV, or if boundaries were supported only by single breakends, the event was classified as **unknown**.

Remaining HLAMP regions were typically embedded within complex genomic rearrangement (CGR) clusters that did not meet the chromothripsis criteria. Within these, we additionally assigned an FBI class to clusters with ≥10 SVs in which FBI were the most frequent SV type. The remaining CGR-associated events were classified as **CGR-TRA** when HLAMP borders were formed by interchromosomal translocations, and as **CGR-NOS** when no boundary translocations were detected.

#### HLAMP Classifier (eicicle)

In this section, we describe eicicle, the algorithm we used to classify HLAMPs as either extrachromosomal (ecDNA) or intrachromosomal (ICamp). It is based on the observation that the copy number consistent with asymmetrical segregation of ecDNAs follows a broad distribution, while the symmetric segregation of ICamps produces a more peaked pattern with copy numbers clustered around a limited number of modes. For computational expediency, we first apply a Log-transformation to the copy number data. In the Log-space, a Gaussian is equivalent to a Log-Normal distribution capable of handling wide data. Our method is based on a structured Gaussian mixture model with two one-dimensional components, where the mixture weights and the mean and standard deviation parameters of each component are parameters of interest. To test a given locus in a given case, our method takes as input the copy number values of all cells at that locus, and outputs the estimated model parameters. We then aggregate these parameters to compute two summary statistics per case: (a) an ecDNA score that is higher for ecDNA cases, and lower for ICamp cases, and (b) a mass-in-window score that is higher in ICamp cases and lower in ecDNA cases. Finally, a classifier based on Linear Discriminant Analysis (LDA) is trained on those cases with the most extreme values (top and bottom 5% quantile for each score) to find a decision boundary which is applied to the remaining cases. In the following, we set up the notation, describe the model, and details of inference.

##### Notation

Let *x*_1_, … , *x_N_* ∈ ℝ_≥0_ denote the observed data. We consider a mixture of 𝑀 = 2 univariate Gaussians with means µ_0_, µ_1_ ∈ ℝ , standard deviations σ_0_, σ_1_ > 0 , and mixing proportions 𝜋 = (𝜋_0_, 𝜋_1_) with 𝜋*_k_* ≥ 0 and 𝜋_0_ + 𝜋_1_ = 1.

##### Model

We consider two scenarios: either the distribution of the observed copy numbers is explained by a wide component, or by two well-separated narrow components. The former corresponds to a majority-ecDNA distribution, while the latter to a majority-ICamp distribution. To mitigate identifiability issues, we introduce a few constraints. To identify the means, we define 𝜇_0_ as a function of 𝜇_0_. To identify the component responsible for the ecDNA, without loss of generality, we assume that if a wide component exists, it is always the first one. To enforce this, we condition the second standard deviation 𝜎_1_ on the first one 𝜎_0_: if 𝜎_0_ < 𝜏 then we constrain 𝜎_1_ ≤ 𝜎_0_, otherwise 𝜎_1_ = 0.1. This creates two distinct regimes: one where two narrow peaks explain the data, and another, where a mostly wide peak explains the data.

##### Generative Model

The probabilistic generative model for eicicle is as follows:

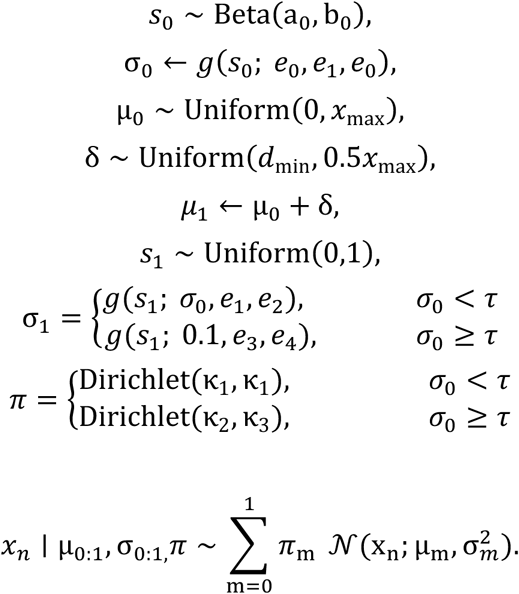

We implemented this model in the probabilistic programming language Pyro^62^ and used stochastic variational inference (SVI)^63^ for inference. SVI is sensitive to initialization, so we repeated model fitting using five random seeds and selected the run with the lowest variational objective. In our implementation, we set *x*_max_ = max *_n_ x_n_*, a_0_ = b_0_ = 0.5, 𝜏 = 0.3, 𝑑_dim_ = 𝜎*_eff_*/2.5, where 𝜎*_eff_* is defined as the robust spread of the data, computed as the difference between the 95^th^ and 5^th^ percentiles, ( 𝜎*_eff_* = 𝑄_0.95_ (*x*) − 𝑄_0.05_(*x*) ). κ_1_ = 5 when 𝜎_0_ < 𝜏 and κ_2_ = 1.0 and κ_3_ = 0.5 when 𝜎_0_ ≥ 𝜏 . We define a clamp function 𝑔(𝑧; 𝑠, ℓ, 𝑢) = min (max (𝑠𝑧, ℓ), 𝑢) and set 𝑒_0_ = *x*_max_/6 , 𝑒_1_ = 10^−3^, 𝑒_2_ = 2, 𝑒_3_ = 10^−4^, 𝑒_4_ = 1.

To summarize each case, we defined two scores: an ecDNA score 𝜓 and a Mass-In-Window metric 𝜂. 𝜓 is defined as a function of the mean and standard deviation of the Gaussian component with the largest 𝜋*_k_*. That is: 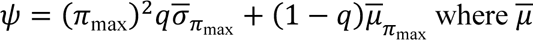 and ̄𝜎 are the standardized mean and standard deviation respectively (across cases), and 𝑞 = 0.66. 𝜂 is defined as the mass in a small neighborhood of size 𝜖 = 0.5 around the mode, that is 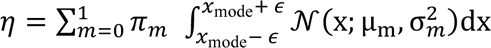

To classify cases into ecDNA vs. ICamp, we use linear discriminant analysis (LDA) using the 5% of extreme cases for training, as implemented in discriminant_analysis. LinearDiscriminantAnalysis function of sklearn library with default parameters. Probability bands at 0.05 and 0.95 are calculated using Platt scaling.

#### ecDNA Simulator

To investigate the dynamics of copy number alterations under ecDNA segregation, we developed a stochastic model. The simulation tracks a population of cells across discrete generations, where the state of each cell 𝑖 at generation 𝑡 is its integer copy number *X_t,i_* ∈ ℕ. Let *N_t_* denote the population size at generation *t*, and let *K* denote the imposed maximum population size (carrying capacity).

##### Population dynamics

Population growth is implemented through a density dependent division probability 𝜆(𝑁_F_) that decreases as the population approaches capacity. In the log-scaled density-dependent division mode used here, each surviving cell divides independently with probability

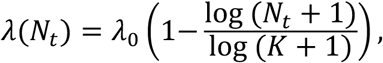

where 𝜆_$_ is the base division probability.

Selection is modeled via a copy number dependent death probability 𝜇(*x*) that induces a U-shaped fitness landscape. Specifically, we apply piecewise constant multipliers to a base death probability

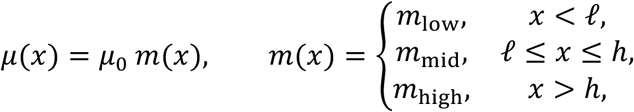

where ℓand ℎ are lower and upper copy number thresholds and u𝑚_HIJ_, 𝑚_4:K_, 𝑚_L:ML_v are scenario specific multipliers. In addition, cells with *x* < 2 are deterministically removed.

##### Copy number dynamics

Each generation consists of three stochastic steps applied to the current population:

1. **Death step.** For each cell with copy number *x*, we sample death as a Bernoulli event with probability 𝜇(*x*). Cells that die or have *x* < 2 are removed from the population.
2. **Division decision.** Each surviving cell divides as a Bernoulli event with probability 𝜆(𝑁_F_).
3. **Copy number update on division.** For each dividing cell with copy number *x*, we first duplicate the copy number (*x* → 2*x*) to mimic DNA replication. These duplicated copies are then partitioned between two daughters via a binomial split:

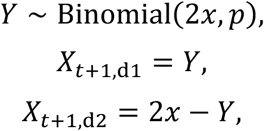

where 𝑝 ∈ [0,1] controls segregation bias (𝑝 = 0.5 is unbiased; 𝑝 > 0.5 biases inheritance toward daughter 1). The two daughters are added to the next generation population. This procedure defines a discrete time, density-regulated branching process with copy number dependent selection and stochastic segregation driven copy number variation. Symmetric division is implemented as a deterministic segregation of DNA material equally between the two daughter cells. Under this condition, the copy number of a clonal population does not change. In our simulations, we used 𝐾 = 10^N^, 𝜆_0_ = 1.0, 𝜇_0_ = 0.01with thresholds ℓ = 100 and ℎ = 500 and multipliers 𝑚_low_ = 2.0, 𝑚_high_ = 4.0, enforced *x* ≥ 2 viability, followed by 𝑝 = 0.6. 𝑚_mid_ = 0.001 and 𝑚_mid_ = 10.0 for high- and low-fitness scenarios respectively.

#### Extrachromosomal Amplicon De-Mixer (ECADeMix)

Given a single-cell WGS dataset in which specific regions have been identified as likely ecDNA, our goal is to infer a) the copy number profiles of distinct ecDNA species present (i.e., *amplicons*) and b) the multiplicity of each species in each cell (i.e., amplicon *loadings*). To solve this problem, we developed a coordinate ascent algorithm using a mixed integer linear program (MILP) to alternatively fix one of these quantities and infer the other, starting by fixing the initial amplicons.

The input to ECADeMix is single-cell CN profiles in 10kb bins, limited to specific regions of interest. To obtain this input, we normalized 10kb read counts using the constant of proportionality between read counts and copy number (i.e., the number of reads per copy per base) from 500kb-based HMMCopy results. Specifically, we took the average number of reads per copy in 500kb bins, divided this number by 50 (to account for the difference in bin size), then divided the 10kb read counts by this value to obtain approximate 10kb copy numbers. We normalized these profiles using non-amplicon-containing cells from the same sample (when available) to account for bin-specific read count biases.

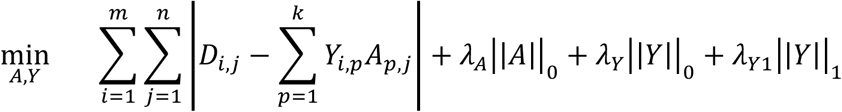

s.t.

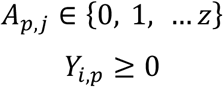

for given number *n* of cells, number *m* of bins, number *k* of amplicons, maximum amplicon copy number *z*, and regularization parameters 𝜆*_A_*, 𝜆*_Y_*, and 𝜆*_Y1_*.

To initialize the MILP, we used either non-negative matrix factorization (scikit-learn implementation) or manual analysis of single-cell CN profiles to generate an initial set of amplicons. The number of amplicons, maximum amplicon CN, and regularization parameters were tuned on a case-by-case basis by inspecting residuals to select the minimal values of *k* and *z* that sufficiently explain the observed data. The parameters (and initialization files where relevant) used for each patient are available in the ECADeMix section of the paper repository (hlamp/pipelines/ECADeMix/metadata). We solve this MILP using the Gurobi optimizer.

#### Calling CN Peaks

To identify CN peaks, i.e., regions where cells cluster around a copy number mode, we used a histogram-based approach followed by 1D smoothing and local-maximum calling. Peaks were defined as local maxima in a one-dimensional signal, subject to user-specified constraints on peak height, prominence, separation, and width. For each locus-specific vector of per-cell copy-number values, we first constructed a 1D histogram (100 bins). The resulting bin counts were smoothed with a Gaussian kernel (scipy.ndimage.gaussian_filter1d, σ = 1.5), and peaks were identified as local maxima in the smoothed histogram using the find_peaks function (scipy.signal) with minimum peak height 1.5, minimum prominence 1.5, minimum peak distance 1 bin, and minimum peak width of 2 bins.

To test whether clones were preferentially enriched in a single peak (rather than distributed across multiple peaks), we computed a clone concentration metric per HLAMP region. For each clone, we computed the fraction of its cells in its most frequent peak and averaged this quantity over clones. For each case, we randomly permuted peak labels across cells within the dataset while preserving clone sizes and peak totals and recomputing the concentration metric to form a null distribution (n = 3,000 permutations). Finally, we combined the one-sided p-values across datasets using Stouffer’s inverse-normal method (as implemented in scipy.stats.combine_pvalues, method=“stouffer”), yielding a single global significance estimate for preferential single-peak enrichment. Prior to Stouffer transformation, we clipped per-dataset permutation p-values away from 0 and 1 to avoid infinite Z-scores.

#### Subclone CN Comparisons

For each ICamp region, we compared subclone-level copy-number patterns across three intervals: i) the full amplicon, ii) the peak (maximum copy) segment within the amplicon, and iii) the interval from the amplicon to the centromere on the same chromosome arm. Using signals package, we computed subclone-level consensus copy number for each interval. Differences in consensus copy number between subclones were evaluated by Student’s *t* test on high-quality cancer cells.

To investigate the mechanisms of ICamp modulation between subclones, we quantified copy-number similarity between subclones by computing the cosine similarity between their copy-number profiles and derived a copy-number similarity score, based on the cosine similarity of the subclone with lower weighted mean relative to the copy-number difference vector between the two subclones.

To differentiate numeric vs. structural modulation of ICamp, we applied a Gaussian mixture model with two components to the copy number similarity score (using GaussianMixture in sklearn.mixture). Datapoints with predicted probability over 0.95 where annotated with their respective class labels: those belonging to the class with the smaller mean CN similarity score were designated as Numeric and the other as Structural. Datapoints with predicted probability less than 0.95 are annotated as Intermediate.

#### Copy Number Correlation between Segments

Copy number of segments that belong to ecDNA were pairwisely compared using Pearson’s correlation, after removing cells with top and bottom 2.5% of the copy-number values to stabilize the correlation. Then, hierarchical clustering was applied to identify the segments where the copy number co-varies. The result was plotted in heatmap using ComplexHeatmap package in R (https://jokergoo.github.io/ComplexHeatmap/).

#### Amplicon Architect

We ran AmpliconSuite-pipeline v1.3.5^64^ for all available scWGS data for each library. We complied result table from Amplicon Classifier function in each case to assess suggested mechanisms of amplification. For case-level analysis, we considered ecDNA-positive when any of the amplicon (from any library) was classified as ecDNA. For region-based analysis, we annotated our scWGS-based HLAMP regions with Amplicon Classifier results, based on their four mechanistic categories (linear, BFB, complex-non-cyclic, ecDNA). To quantify the agreement between Amplicon Architect-based method and single cell copy-number distribution-based classification, we calculated Cohen’s kappa using psych package in R.

### scRNA-seq Analyses

#### Preprocessing

All analyses for single cell RNA sequencing data were performed in Python (v3.10.16) using Scanpy (v1.10.4)^65^. FASTQ sequencing files were aligned to the GRCh38 reference genome, barcodes were filtered, and unique molecular identifiers (UMIs) were identified using the 10x Cell Ranger software (v6.0.1). Cells with fewer than 3 genes or with more than 40% mitochondrial content were removed. Highly variable genes were selected using the highly_variable_genes function with the flavor option seurat_v3. Raw counts were then normalized by library size and log-transformed with a pseudo-count of 1. Principal component analysis (PCA) was performed retaining the top 50 PCs, which were used for Leiden clustering and for generating the UMAP^66^ embedding for visualization.

#### Cell Type Annotation

The cell types were defined using the markers reported in the previous study by our group^67^. Each cell was scored for each of the main cell types using the function score_genes. Then resulting scores were z-score normalized, then averaged per leiden cluster. Each cluster was annotated based on the cell type with the highest score. In turn, each cell inherited its cluster’s annotation.

#### Identifying Malignant Cells

To identify malignant cells in the epithelial compartment, single-cell copy number signal was computed per sample using infercnpy v0.6.0 (https://github.com/icbi-lab/infercnvpy)^68^, using a subset of non-epithelial cells as reference. Copy number clusters were identified by first computing top Principal components, and then community-based clustering.

#### Pathway Analysis

##### CCND1 loss in OV-052

Differential expression was performed for copy number cluster cluster “3” versus all other cells using Scanpy’s Wilcoxon rank sum test using function rank_genes_groups after library size normalization and log transformation (and identification of highly variable genes). Preranked gene list was constructed by sorting genes by signed effect size (log fold change). This ranked list was used as input to GSEApy (v1.1.4)’s preranked GSEA (prerank) against the MSigDB Hallmark gene sets with 1,000 permutations and gene set sizes restricted to 15 to 500 genes. Enrichment results were saved and summarized with a bar plot of the top positively and negatively enriched pathways.

##### Pathway activation in GBM0721

Gene sets for the Hallmarks pathway were acquired from msigdb_v2024^68^. For each hallmark pathway score, a multivariable linear regression model was fit with z scored EGFR and MDM4 expression as predictors, including standard scRNA seq QC covariates (total counts, detected genes, mitochondrial fraction) and an interaction term. Specifically, for each score 𝑦, the model 𝑦 ∼ *x*_!_ + *x*_+_ + *x*_!_*x*_+_ + QC was estimated by ordinary least squares after removing cells with missing values, and scores with fewer than 10 complete observations were excluded. Benjamini–Hochberg correction was applied across pathway scores separately for each coefficient. Pathways were flagged as showing a concordant joint association when both *x*_!_ and *x*_+_ were significant after FDR correction (𝑞 ≤ 0.05) and their coefficients had the same sign.

#### Enrichment and Depletion of co-expression Patterns in scRNAseq

For each dataset and gene pair, we first defined per-cell “overexpression” using raw UMI counts and a depth-adjusted Negative Binomial (NB) model. For each gene 𝑔 in cell 𝑖, the expected mean count 𝜇_GS_ was set proportional to library size total_counts_G_, and gene-specific dispersion 𝜃_S_ was estimated under an NB2 variance parameterization Var(𝑦_GS_) = 𝜇_GS_ + 𝜇^+^ /𝜃_S_. We then computed an NB Pearson residual

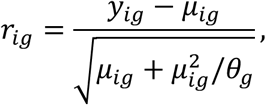

and called a gene “high” in a cell if 𝑟_GS_ > 2. Cells were assigned to one of four categories for each pair (EGFR-only high, partner-only high, both high, neither), yielding a 2 × 2contingency table 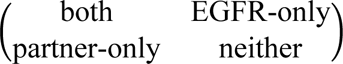. We tested for association between “EGFR-high” and “partner-high” using Fisher’s exact test, with a one-sided alternative for depletion (*EGFR*–*CDK4*) or enrichment (*EGFR*–*MDM4*), and report odds ratios and p-values as above.

#### Fluorescence in situ Hybridization

Cell lines were cultured in their respective media and arrested in metaphase with KaryoMAX^TM^ Colcemid^TM^ Solution (Thermo Fisher Scientific, #15212012). Colcemid conditions were 0.05 μg/mL for 2 hours (NCI-H524, NCI-H69, NCI-H82), 0.05 μg/mL for 6 hours (COLO320-DM, COLO320-HSR), 0.05 μg/mL for 6 hours to overnight (GBM39-DM, GBM39-HSR), or 0.10 μg/mL for 2 hours (2765_2). Cells were harvested as a single-cell suspension, incubated in 75 mM KCl for 10 minutes at 37℃, and fixed in ice-cold methanol:glacial acetic acid (3:1; Carnoy’s fixative) at −20℃ for 30 minutes to overnight. Cell pellets were washed 3 additional times with Carnoy’s fixative and dropped onto glass slides to generate metaphase spreads.

Slides were processed for fluorescence *in situ* hybridization (FISH) through graded ethanol (70%, 85%, and 100%) and air-dried. FISH probes were either purchased (e.g., *MYC*, MetaSystems #D-6008-100-OG; *MYCN*, Empire Genomics SKU *MYCN*, CHR02) or synthesized in-house from BAC clones (*Mdm2* RP23-428D5; *Cen10* RP23-309H16; *EGFR* RP11-433C10 and RP11-339F13; *CEN7* p7t1). For COLO320-DM/COLO320-HSR lines, DNA paint probes (MetaSystems) were applied to slides, sealed with a coverslip using rubber cement, co-denatured with the specimen at 72℃ for 3 minutes, and hybridized overnight at 37℃ in a humidified chamber. Post-hybridization, slides were washed in 2X SSC three times for 2 minutes each, rinsed in PBS, counterstained with DAPI, dehydrated through graded ethanol (70%, 85%, and 100%), and mounted in ProLong Gold antifade (all cells except COLO320 lines; #P36930) or ProLong Diamond antifade (for COLO320-DM/COLO320-HSR; Thermo Fisher, #P36965).

Images were acquired either on a Zeiss LSM880 confocal microscope (Carl Zeiss Microscopy) using a Zeiss 63X/1.41 NA oil immersion objective (COLO320-DM/COLO320-HSR) or on an ECHO Revolve microscope (Discover Echo) using a 60x oil objective (all other cell lines).

---

**Extended Data Fig. 1.**
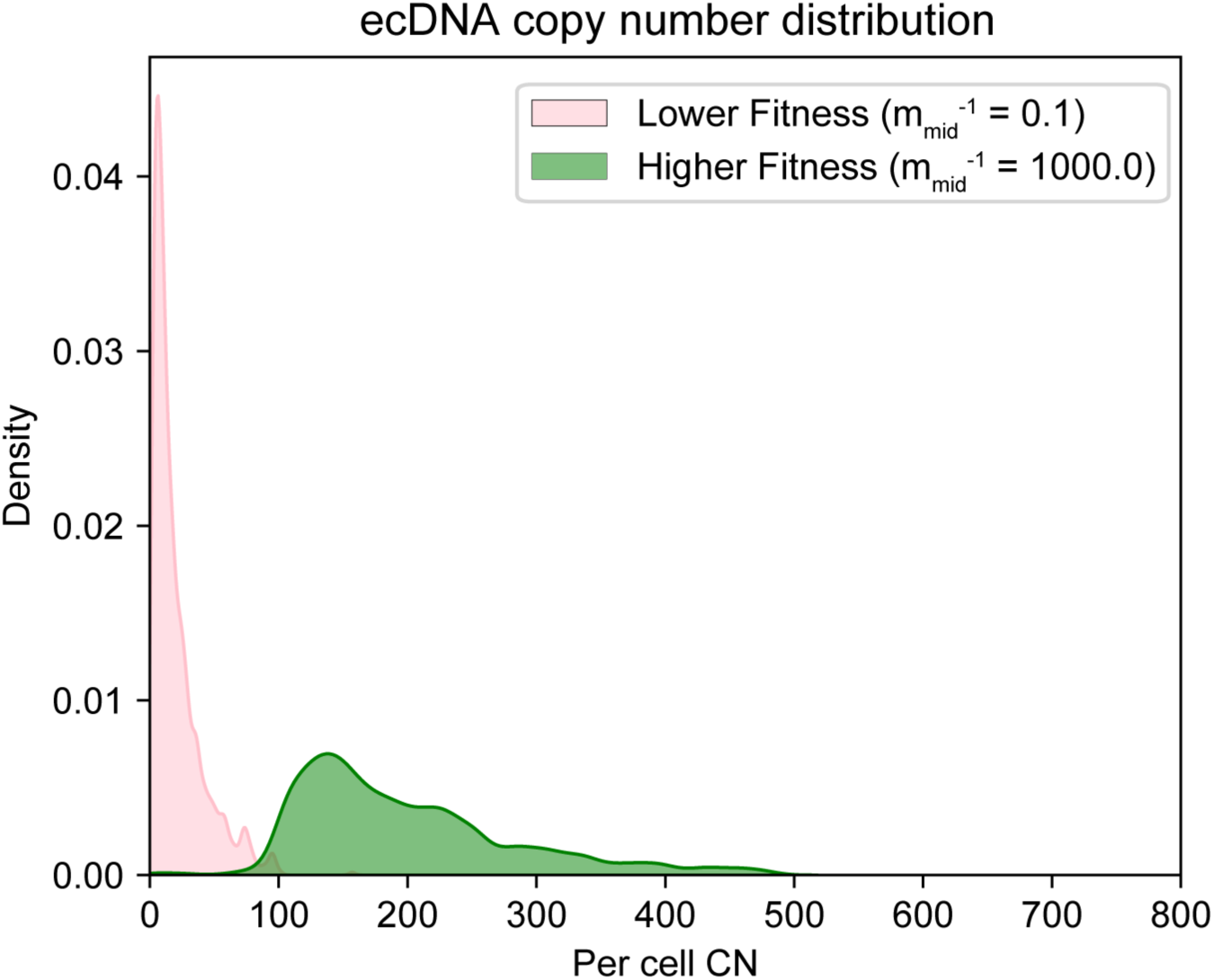
Probabilistic simulation model for high-level copy-number amplification. Copy-number histograms generated from simulations of ecDNA segregation after 500 generations; 1,000 cells were randomly subsampled to mimic sequencing. The green population has higher net fitness than the pink population. In this model, cells with copy number >500 incur a twofold higher fitness cost than cells with copy number <100. Net fitness for cells with copy number 100-500 is shown in the legend (**Methods**).

**Extended Data Fig. 2.**
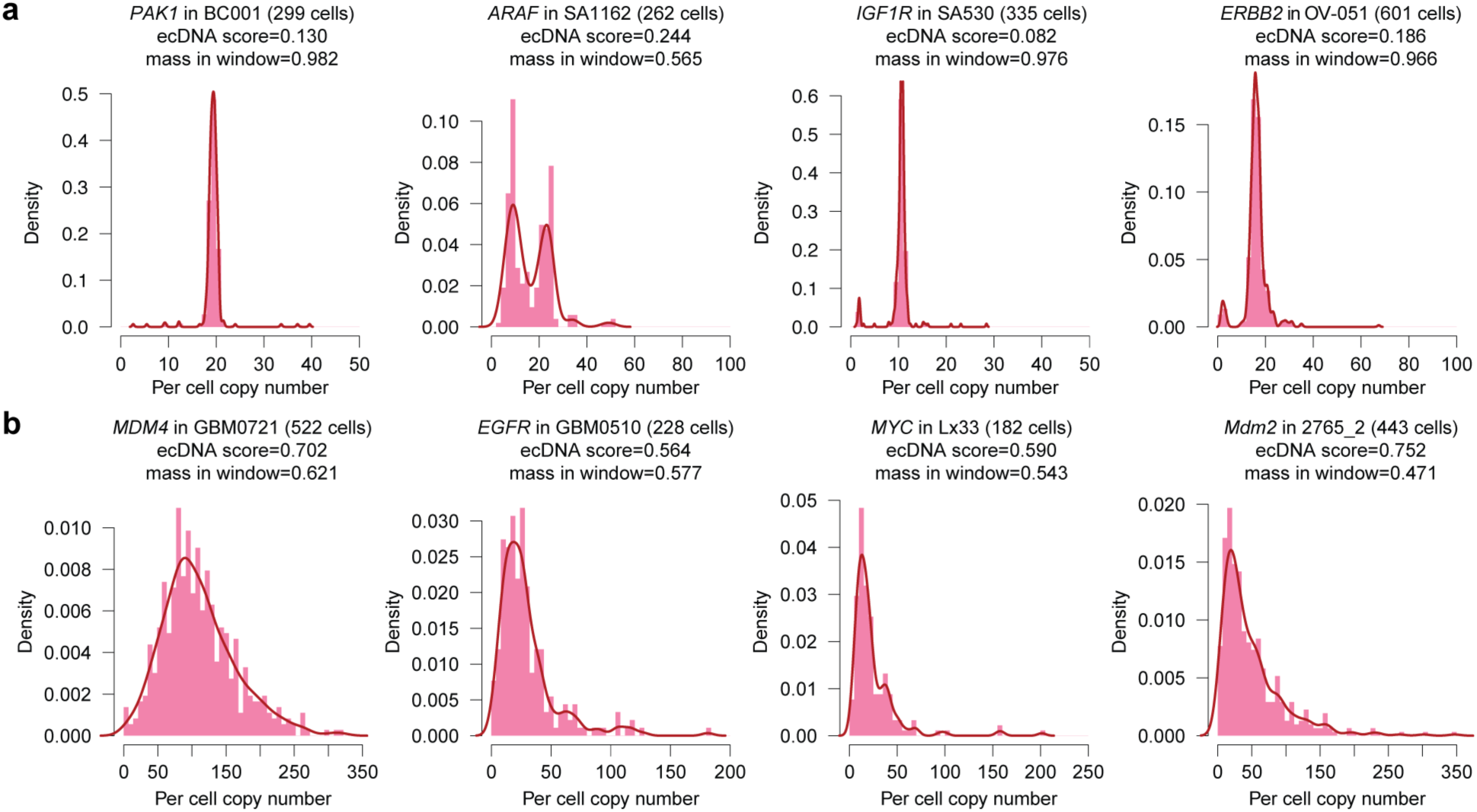
Distinct single-cell copy-number distributions for ecDNA and ICamp. **a-b**, Copy-number histograms (pink bars) and corresponding density estimates (red line) for HLAMP regions harboring *bona fide* oncogenes in 4 cases with ICamp **(a)** and 4 ecDNA cases **(b)**. Density plots were computed by kernel density estimation in base R.

**Extended Data Fig. 3.**
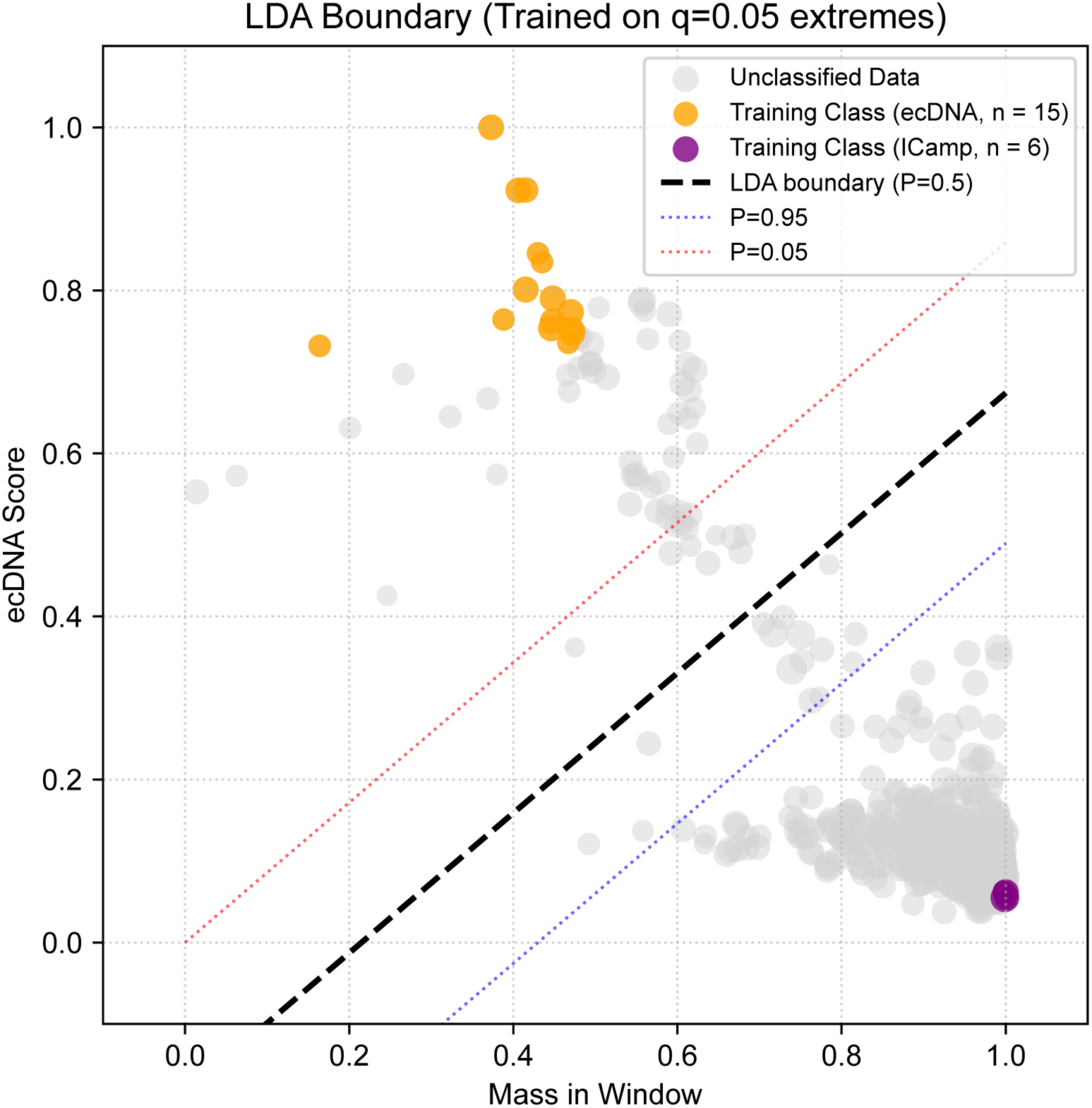
Machine learning-based differentiation of ecDNA from ICamp using. eicicle. A probabilistic model scores each HLAMP region according to the properties of its single-cell copy number distribution. A linear classifier, trained on the extreme values of these scores, defines a decision boundary between ecDNA-positive and ecDNA-negative regions (**Methods**). Red and blue dotted lines show the 0.95 and 0.05 probability bands, respectively. LDA; linear discriminant analysis.

**Extended Data Fig. 4.**
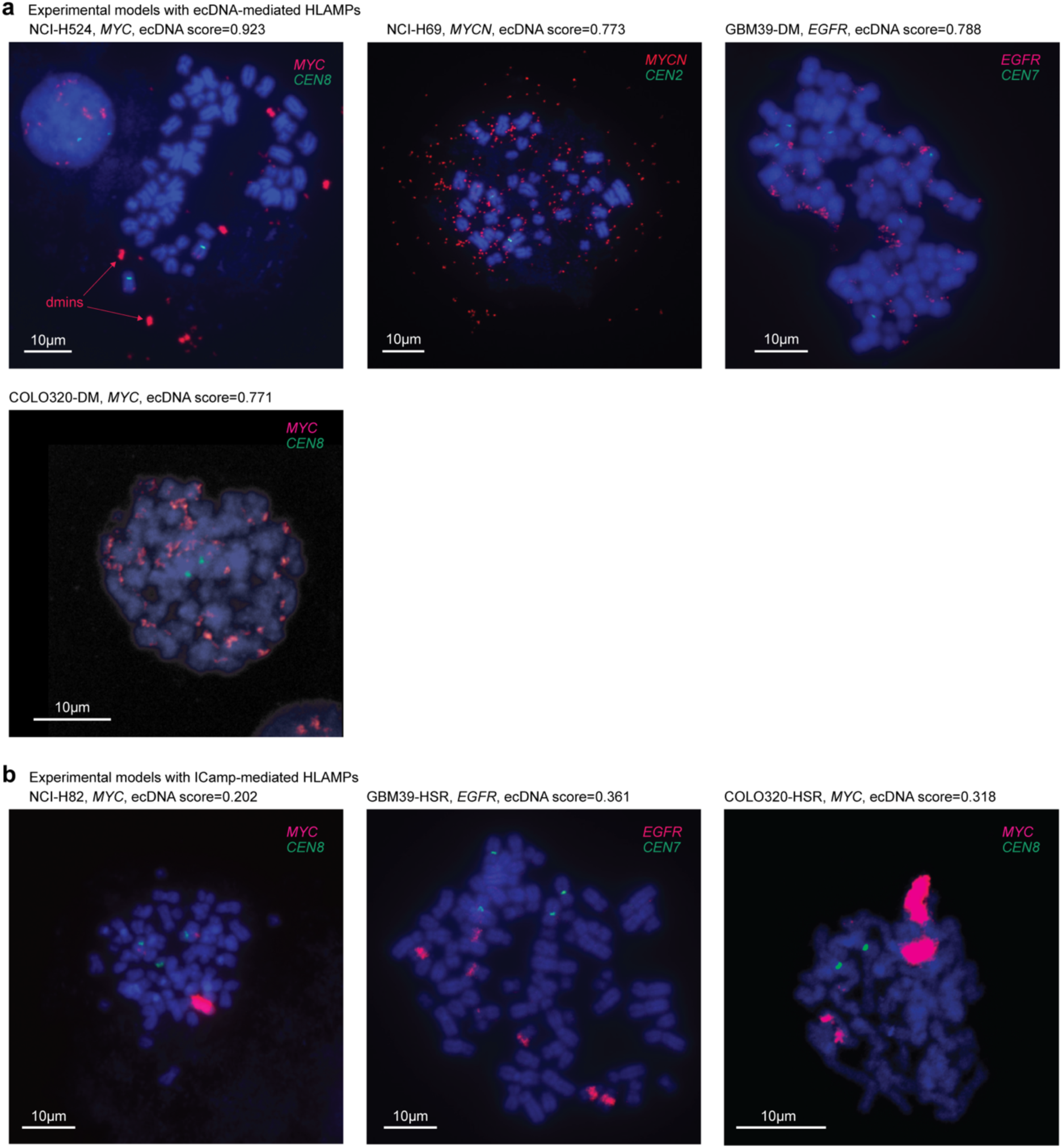
Cytogenetic features of HLAMP regions in experimental models. **a-c**, Cytogenetic evidence for oncogene amplification assessed by metaphase DNA fluorescence *in situ* hybridization (FISH). **a**, NCI-H524 (**top left**) is a small-cell lung cancer (SCLC) suspension cell line with high ecDNA score. The *MYC* probe (red) highlights a compact, intensely fluorescent extrachromosomal structure with bilobed morphology, consistent with double minutes (dmin). NCI-H69 (**top middle**) is a SCLC cell line grown in suspension. *MYCN* FISH shows numerous small, dispersed extrachromosomal signals consistent with *MYCN*-bearing extrachromosomal circular DNA (ecDNA). GBM39-DM (**top right**) and COLO320-DM (**bottom left**) show ecDNA-mediated amplification of *EGFR* and *MYC*, respectively. Consistent with these findings, these models had high ecDNA scores in our single-cell copy-number (CN) distribution analysis using eicicle. **b**, NCI-H82 (**left**) is an SCLC cell line with mix adherent and suspension growth. The *MYC* probe reveals intrachromosomal clustered amplification as a homogeneously staining region (HSR), with no ecDNA detected. GBM39-HSR (**middle**) and COLO320-HSR (**right**) show HSR-mediated amplification of *EGFR* and *MYC*, respectively. Consistent with these findings, these models had low ecDNA scores.

**Extended Data Fig. 5.**
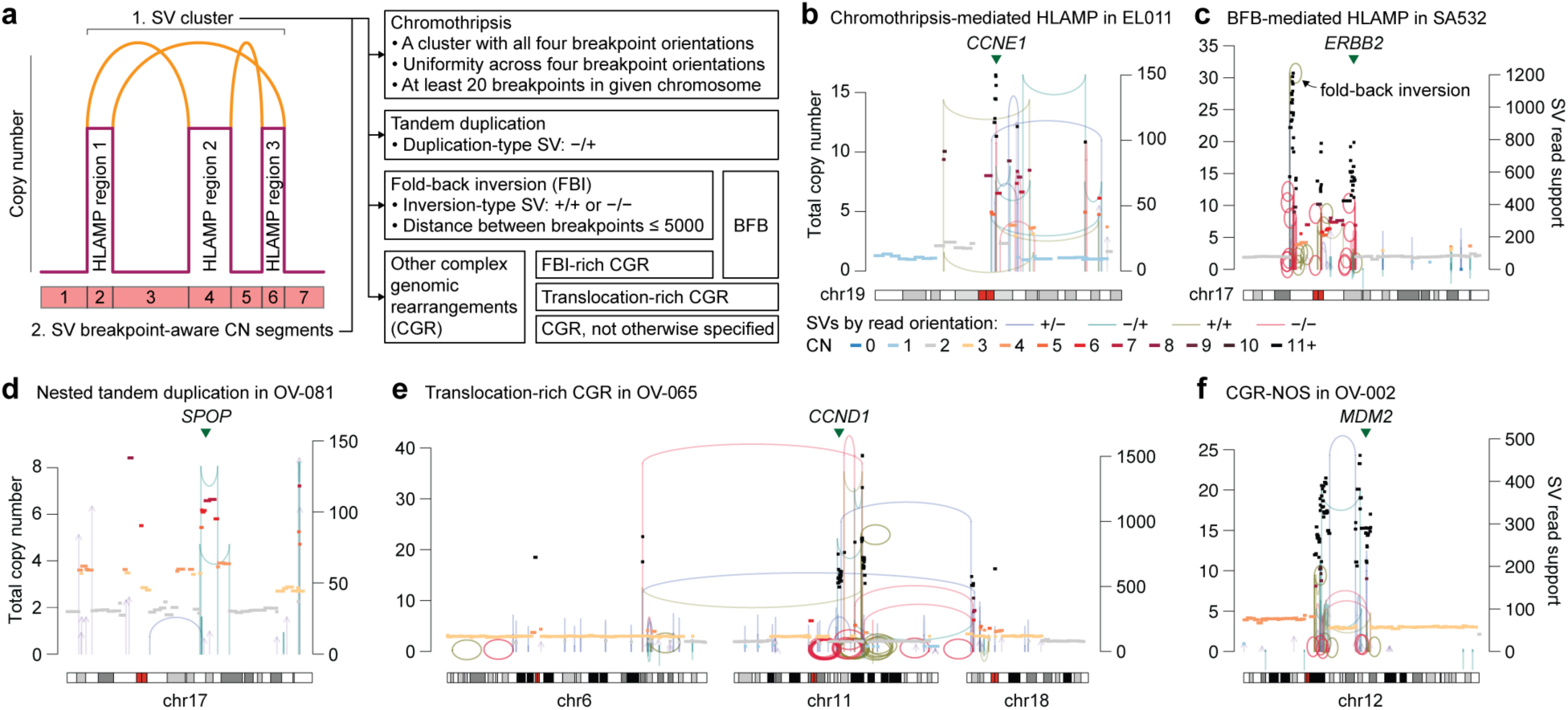
Classification of HLAMP regions by initiating mechanism. **a**, Schematic illustration of HLAMP regions and their relationship to structural variants (SVs). SV breakpoints were clustered into event footprints using clusterSV^23^, accounting for breakpoint proximity, background SV rates and size distributions for each SV type. HLAMP regions were classified into 5 mechanistic categories (and unknown; **Methods**) based on SV features at their boundaries^21^. **b**, A *CCND1* HLAMP event with multiple SVs in all 4 orientations, indicating chromosomal fragmentation and random ligation, consistent with chromothripsis. **c**, A multifocal HLAMP event amplifying *ERBB2*. Major copy-number boundaries exhibit fold-back inversions (FBIs), a genomic footprint suggestive of ligation between sister chromatids. Multiple FBIs facing each other suggest generation by breakage-fusion-bridge (BFB) cycles. **d**, Two overlapping tandem duplications that together quadrupled a segment containing *SPOP* in an ovarian cancer case. **e**, A complex genomic rearrangement amplifying *CCND1*. Interchromosomal translocations at multiple borders between HLAMP segments and unamplified regions suggest that these translocations preceded copy-number amplification. **f**, A complex genomic rearrangement within chromosome 12, amplifying *MDM2*.

**Extended Data Fig. 6.**
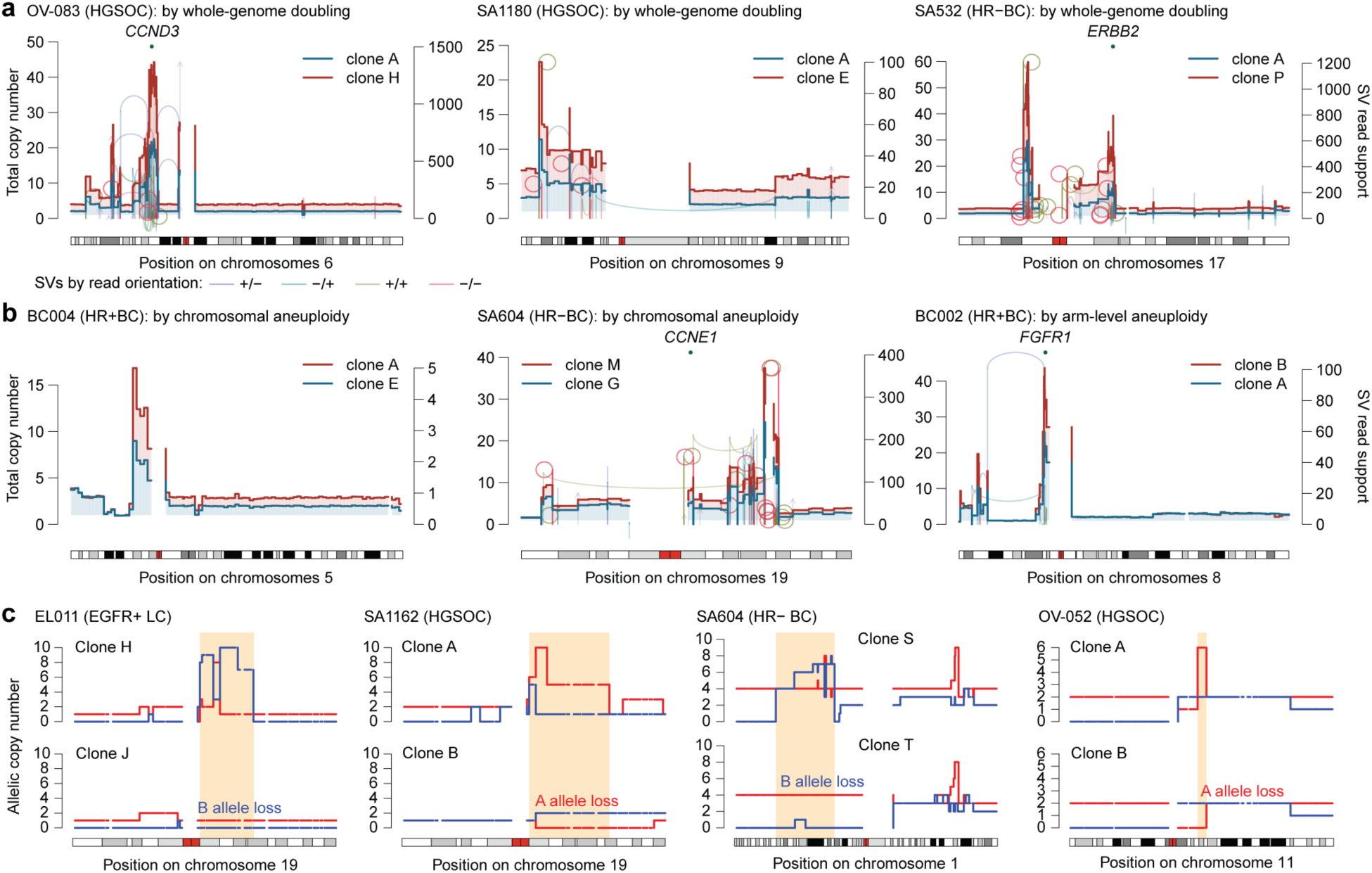
Numeric modulation of intrachromosomal amplification. **a**, Representative examples of subclone-specific duplication of ICamp via whole-genome doubling. In each case, the major clone (clone A; red line) exhibits a diploid-ranged copy number (CN) profile, whereas a minor subclone with whole-genome doubling (turquoise line) shows a global doubling of the CN profile, including the ICamp regions. **b**, Examples of aneuploidy-mediated ICamp duplication. ICamp is further amplified by subclone-specific whole-chromosomal aneuploidy (**left** and **middle** panels) and arm-level aneuploidy (**right** panel). **c**, Four cases showing subclone-specific loss of ICamp due to aneuploidy. Allele-specific CN analysis by the Signals package^17^ reveals loss of the allele carrying the HLAMP event in subclones, resulting in loss of heterozygosity. These findings indicate that the subclone without HLAMP (exhibiting LOH) is not ancestral to the other subclones with HLAMP; instead, it represents a daughter lineage generated by aneuploid loss of the HLAMP-carrying allele. In EL001 and SA1162, the HLAMPs target *CCNE1*, and in OV-052 it targets *CCND1*.

**Extended Data Fig. 7.**
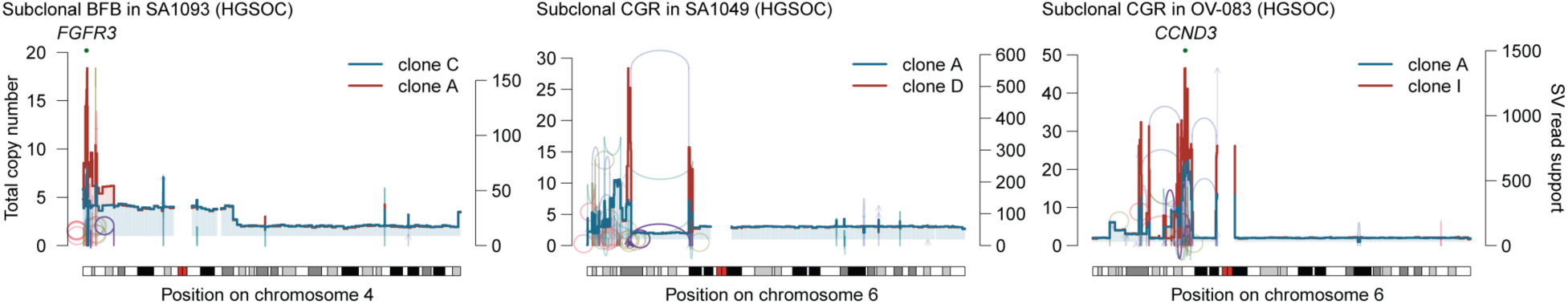
Structural modulation of intrachromosomal amplification. Structural modulation of HLAMP in three representative cases. A subclonal fold-back inversion (purple line) produces a subtelomeric copy-number (CN) gain in chromosome 4, further amplifying *FGFR3* (**left**). Subclonal complex genomic rearrangements (CGR; purple lines) increase CN at HLAMP regions (**middle** and **right**).

**Extended Data Fig. 8.**
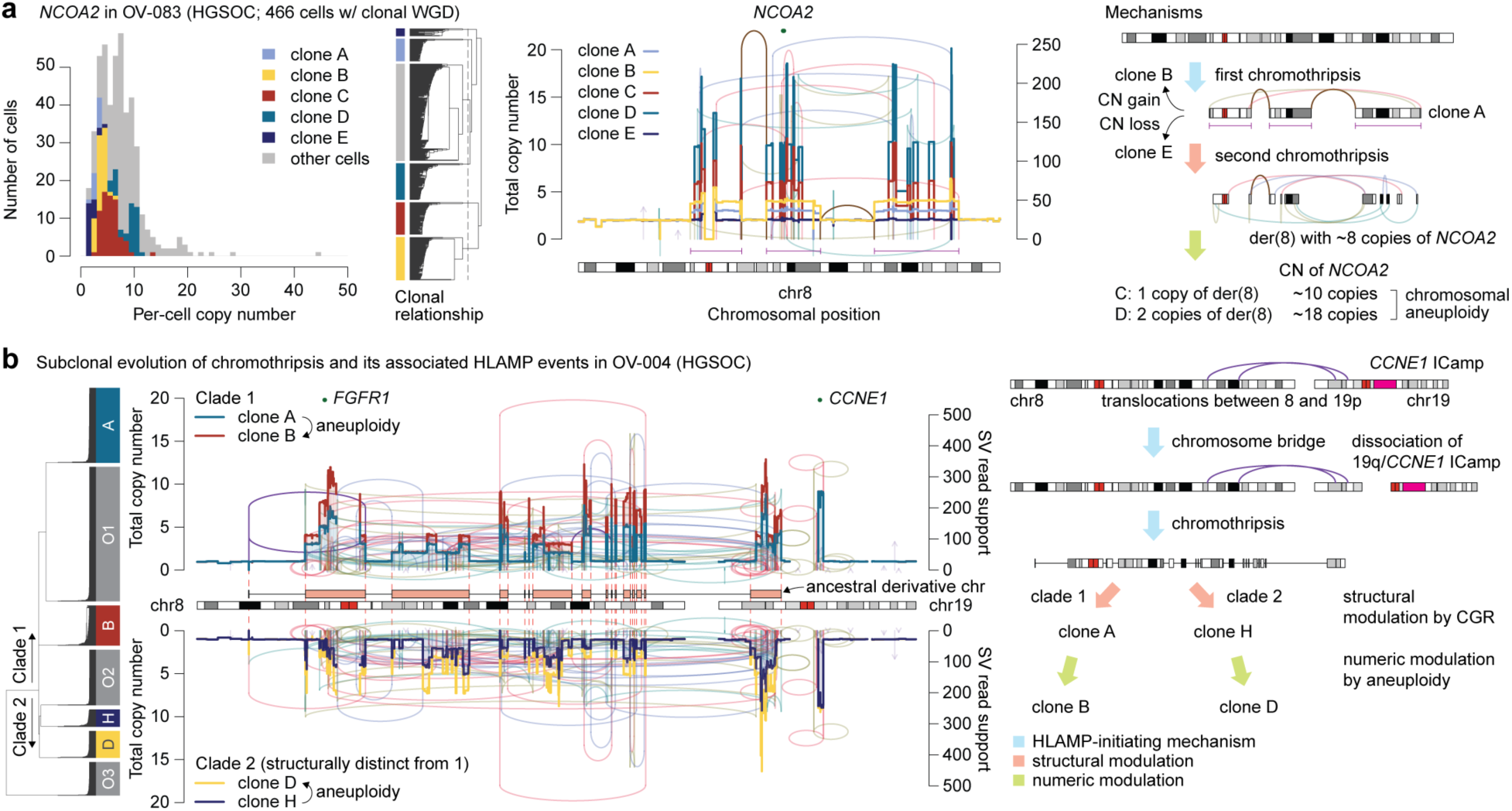
Diversification of HLAMP events after chromothripsis. **a**, An ICamp event involving chromosome 8 in an ovarian cancer case (OV-083), initiated by chromothripsis, followed by a second chromothripsis and subsequent whole-chromosome aneuploidy. **b**, Another ovarian cancer case (OV-004) with chromothripsis and subclone-specific CN profiles for two major phylogenetic clades (**left**). Both clades share the same segmental CN gain, indicating the ancestral structure of the derivative chromosome after chromothripsis (**middle**). This derivative chromosome underwent distinct repair trajectories, leading to divergence into two clades. Within each clade, subclones were further diversified by whole-chromosome duplication. A schematic illustration of the described mechanism (**right**).

**Extended Data Fig. 9.**
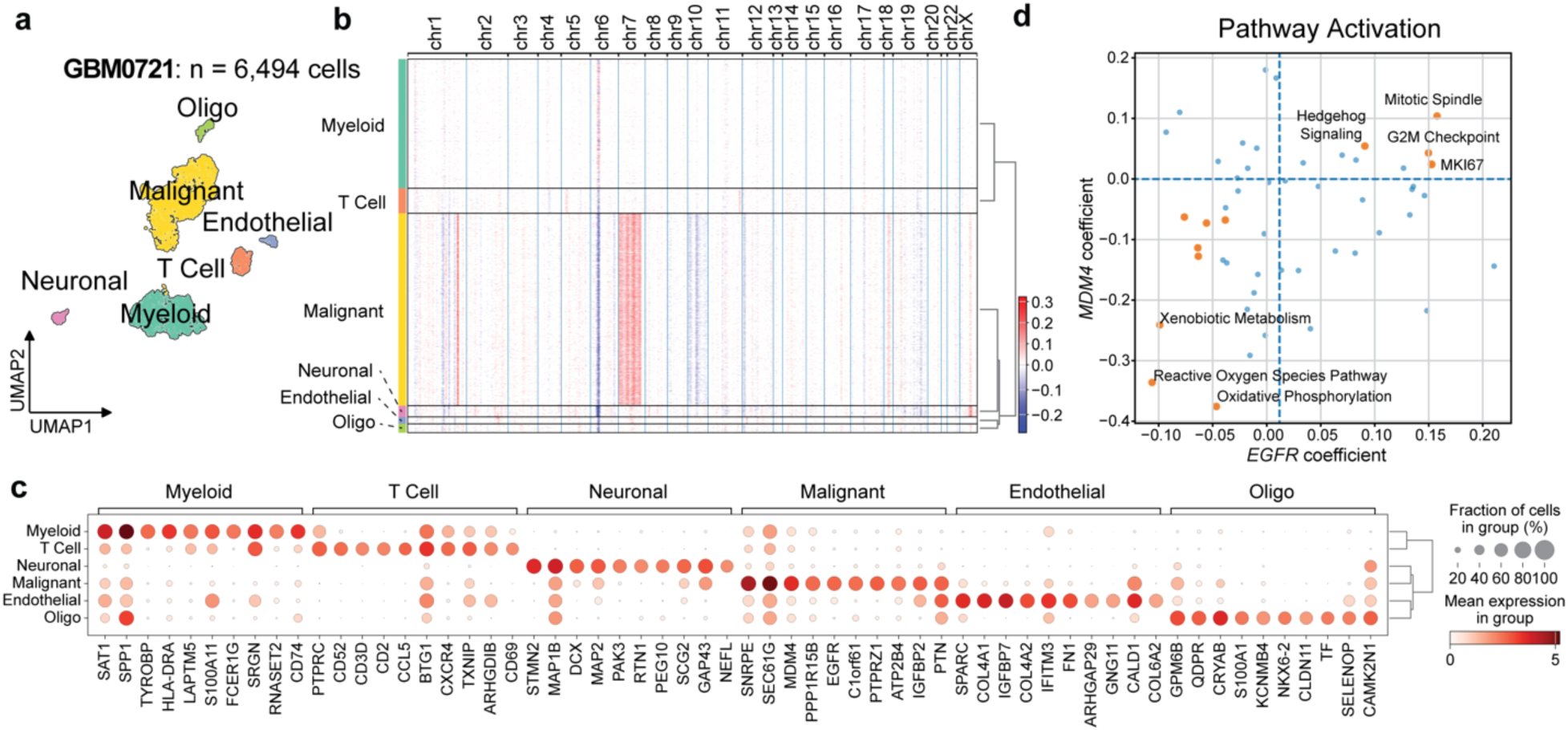
Single-cell RNA-seq analysis in glioblastoma samples. **a-c**, UMAP embedding of 6,494 cells passing quality control from scRNA sequencing of GBM0721 (**a**). Major cellular components are annotated using inferred copy number (CN) signals (**b**) and differentially expressed marker genes (**c**). **d**, Standardized partial regression coefficients linking hallmark pathway scores to expression of *EGFR* and *MDM4* in malignant cells, adjusted for quality metrics (**Methods**).

**Extended Data Fig. 10.**
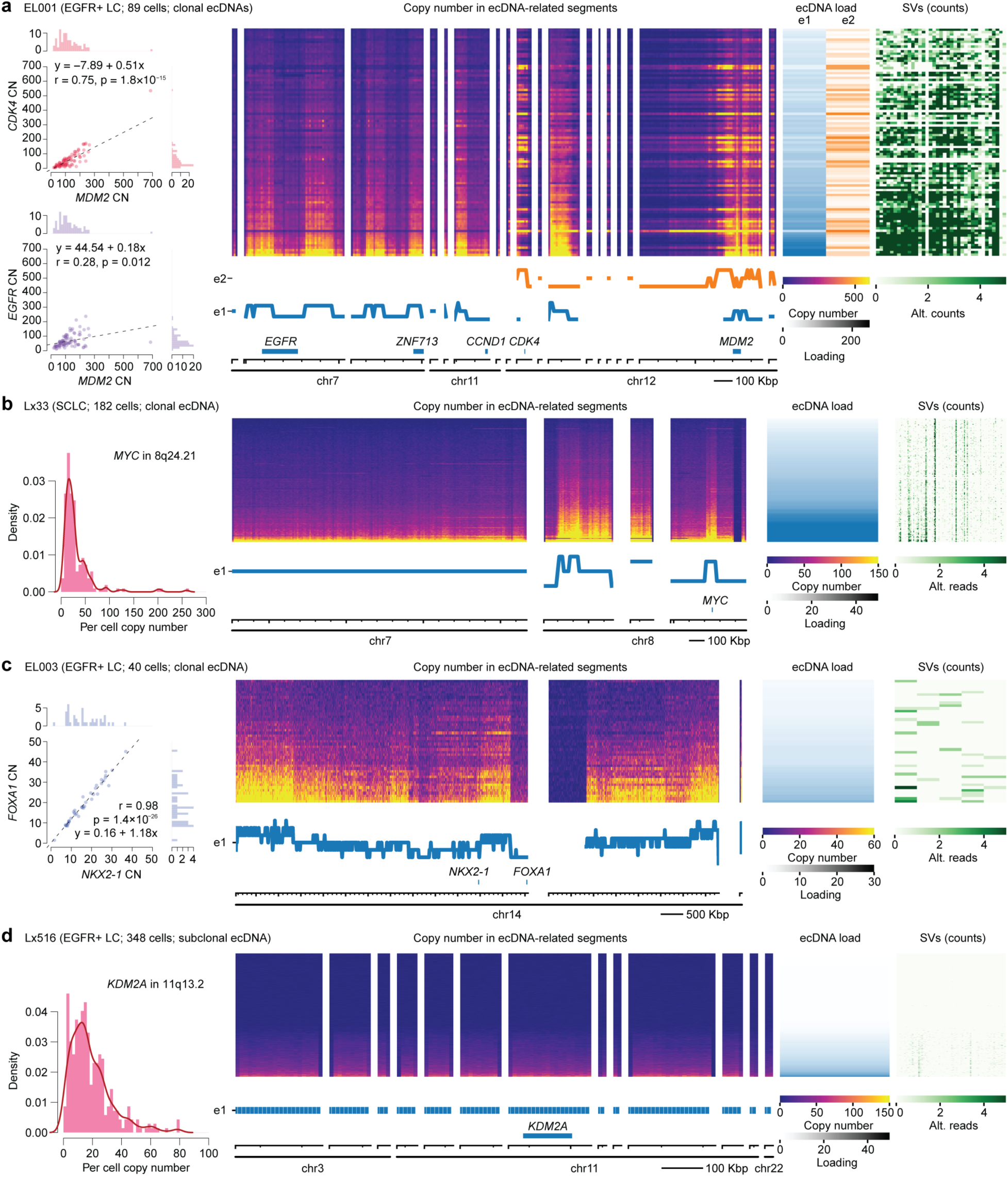
ecDNA architecture revealed by ECADeMix. Decomposition of ecDNA species based on ECADeMix. Shown are four cases in which ecDNA species exhibit minimal structural variation across cells **a-d**. Copy-number (CN) distribution of key oncogenes and their pairwise correlations (**left**), per-cell CN heatmaps and inferred structures of ecDNA species (**middle**), per-cell ecDNA species loads and associated SVs at single-cell resolution (**right**). EL001 was sequenced at greater depth (≥0.1x per-cell coverage), increasing sensitivity for SV detection.

**Extended Data Fig. 11.**
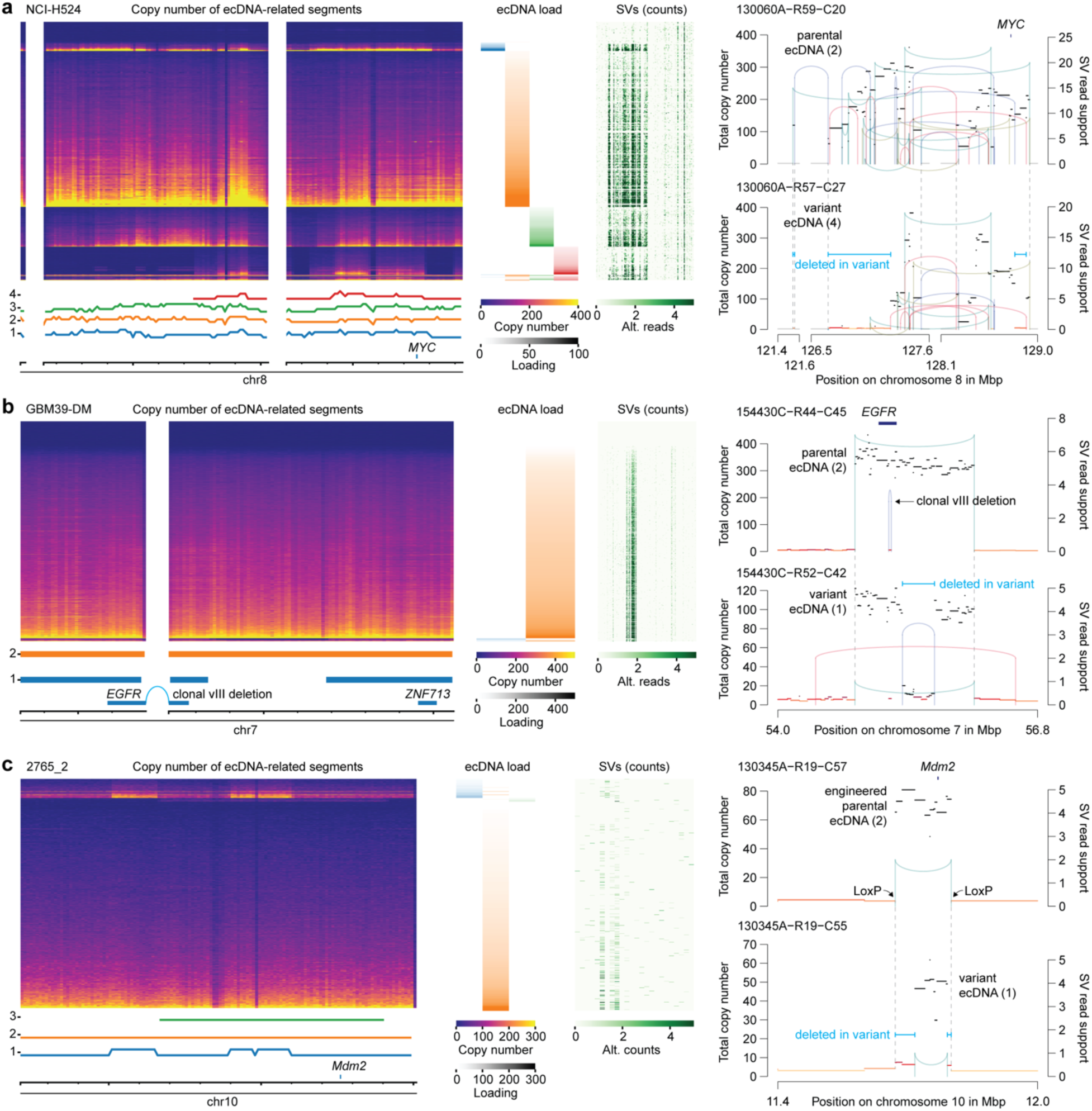
Divergent evolutionary trajectories driven by ecDNA remodeling. **a-c**, Three representative cases in which multiple ecDNA species target the same locus, with evidence of divergent evolution. In each case, one species spans the full set of HLAMP regions, consistent with the parental ecDNA structure (**left**). Variant ecDNA species harbor intramolecular rearrangements that produce segmental deletions (**right**). The SCLC cell line NCI-H524 shows ecDNA targeting *MYC* generated by complex genomic rearrangements **(a)**, GBM39-DM cells harbor *EGFR* ecDNA with the variant III deletion **(b)**, and the genetically engineered mouse model 2765_2 contains engineered ecDNA targeting *Mdm2* **(c)**.

**Extended Data Fig. 12.**
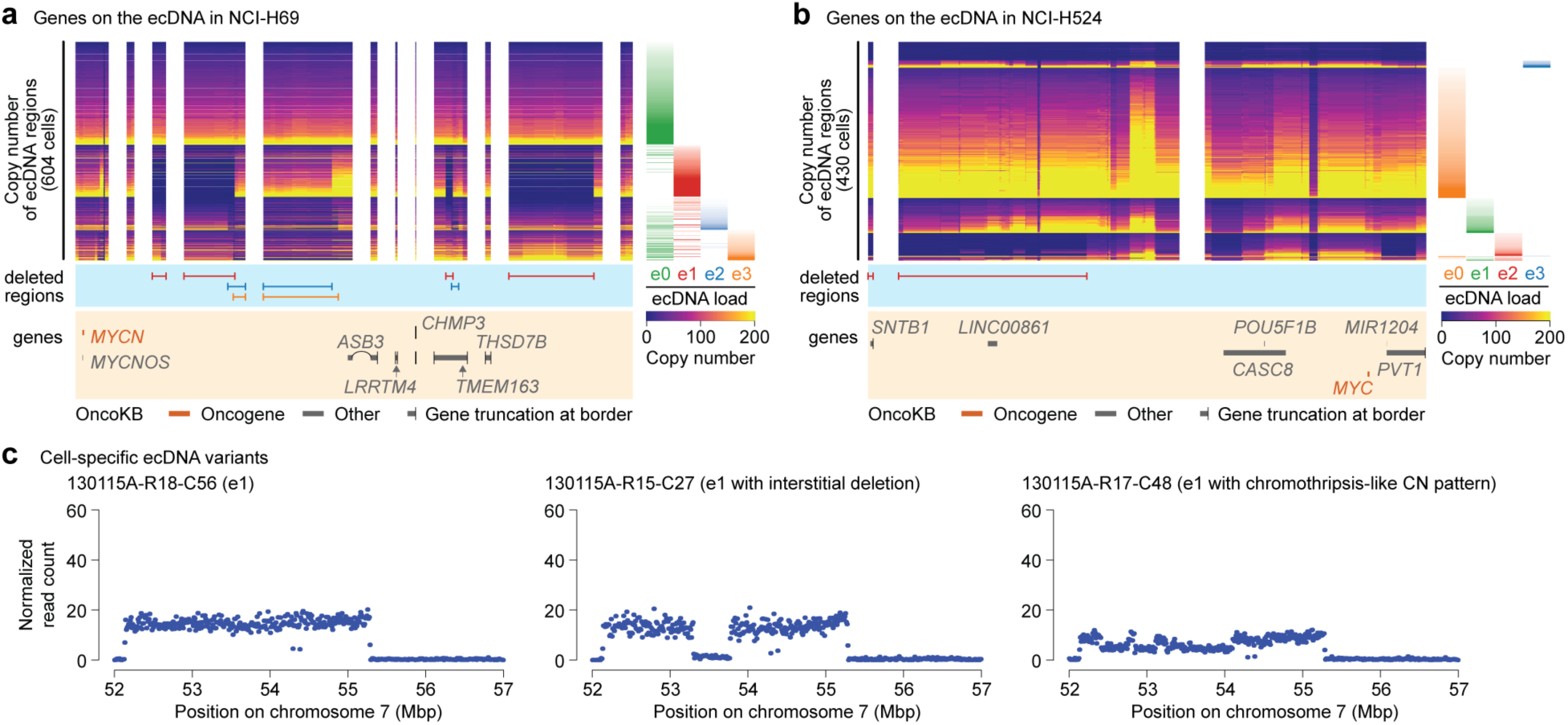
Structural variants underlying divergent ecDNA evolution. **a-b**, Variant ecDNA-specific deletions and their relationships to genes on ecDNA molecules in NCI-H69 **(a)** and NCI-H524 **(b)**, two cases with prevalent ecDNA variants. In both cases, deletions primarily affect gene-poor regions, regions containing already-truncated genes, or long noncoding RNAs. In contrast, *bona fide* oncogenes (*MYCN* in NCI-H69 and *MYC* in NCI-H524) are preserved. **c**, Cell-specific copy-number (CN) patterns indicative of ongoing ecDNA evolution. Shown are an intact e1 ecDNA in GBM0510 (**left**), a variant with a central deletion (**middle**), and a variant with an oscillating CN pattern (**right**), consistent with chromothripsis on ecDNA.

**Extended Data Fig. 13.**
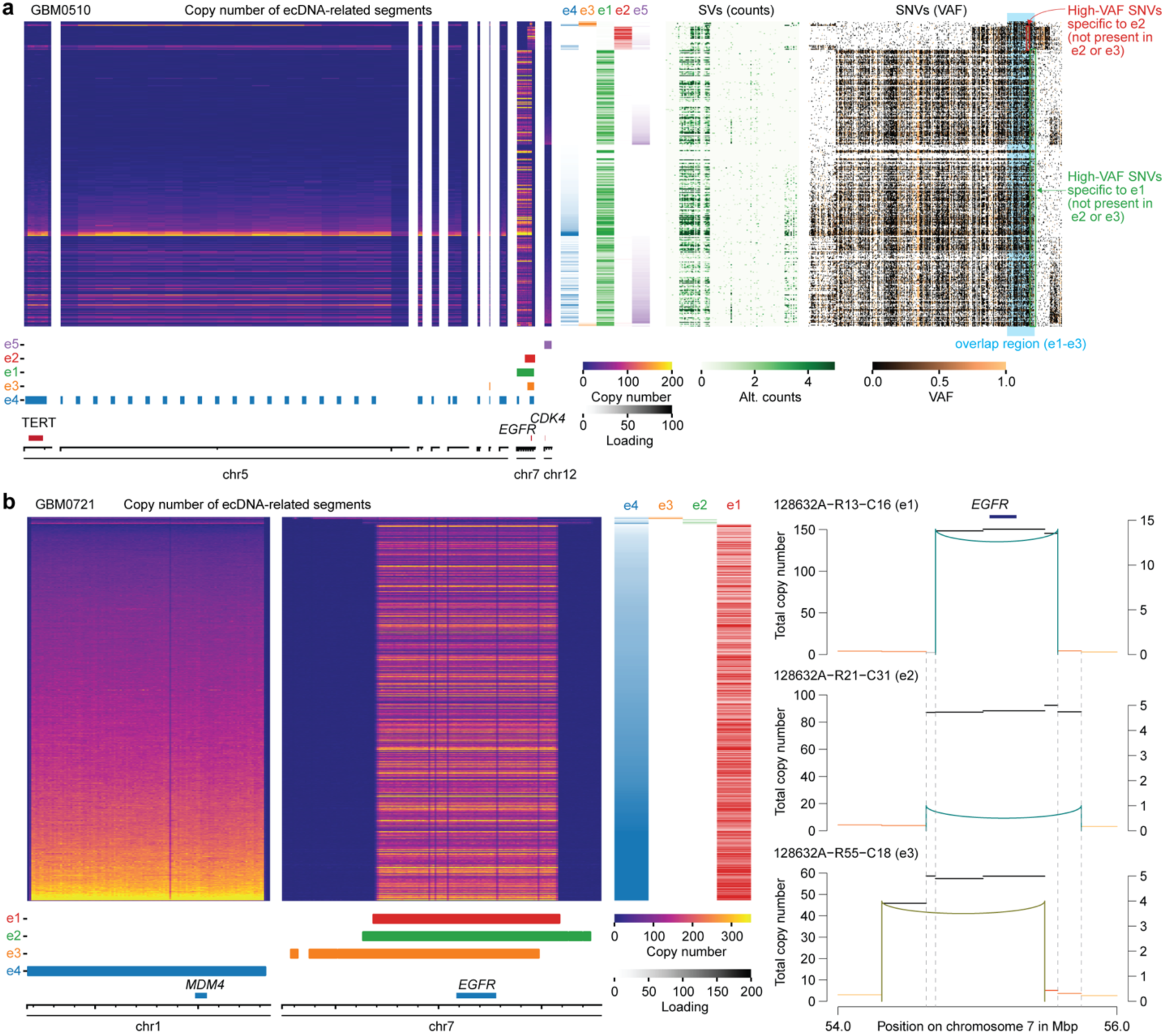
Convergent evolution in glioblastoma via recurrent acquisition of *EGFR* ecDNAs. **a**, GBM0510 shows convergent evolution, with three *EGFR* ecDNA species arising from distinct structural variants (middle heatmap) and defining three mutually exclusive clones. These three clones share many single-nucleotide variants (SNVs) and differ by a small number of clone-specific SNVs (indicated by red and green arrows), consistent with late emergence of *EGFR* ecDNAs during tumor evolution. **b**, GBM0721 similarly harbors three distinct *EGFR* ecDNA species attributable to different structural variants (e1-e3). ecDNA e1 is predominant, whereas e2 and e3 are found in a minority of cells. In contrast, only a single *MDM4* ecDNA species is present, consistent with earlier acquisition. SVs underlying e3 could not be fully resolved because of the small number of e3-positive cells.

**Extended Data Fig. 14.**
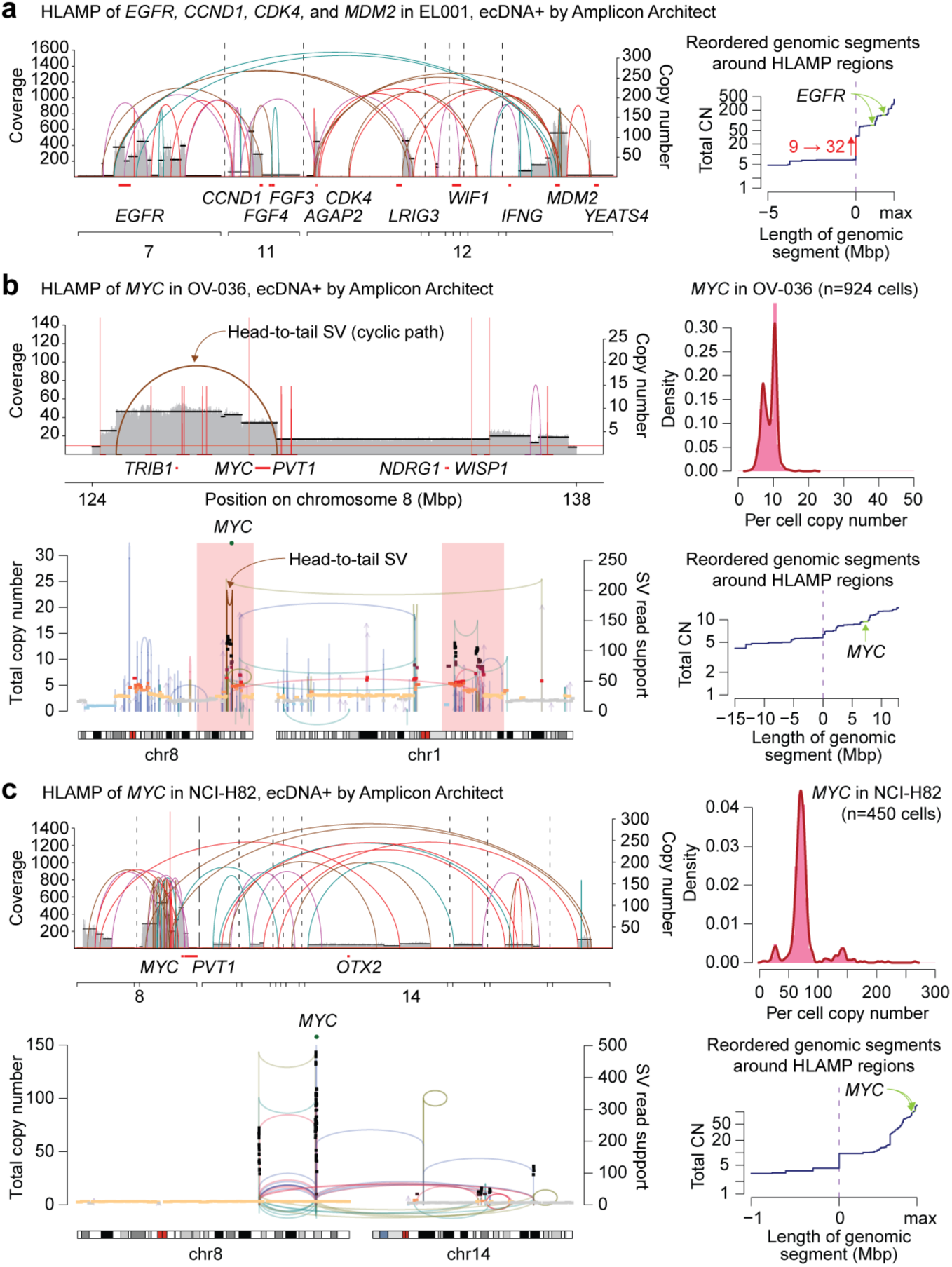
Discrepancies between bulk genome graph-based and single-cell copy number distribution-based ecDNA definitions. **a**, In EL001, Amplicon Architect (AA) classified the amplicon as ecDNA (**left**), consistent with its single-cell copy-number (CN) distribution. These regions exhibit an abrupt CN transition (from 9 to >30 copies; **right**), characteristic of ecDNA-positive cases. However, because of complex connectivity, AA grouped all segments into a single ecDNA, whereas our analysis resolved two independent ecDNAs (Fig. 3). **b**, In OV-036, AA identified a cyclic genome graph around the *MYC* locus (**top left**) and classified it as ecDNA. However, the single-cell CN distribution for *MYC* shows 2 prominent peaks (**top right**), consistent with an ICamp with intratumoral heterogeneity. At the pseudobulk level, the *MYC* amplicon is densely connected by complex genomic rearrangements to regions on chromosome 1 (**bottom left**). These segments (red shading) show a gradual, stepwise increase in CN toward the *MYC* locus (**bottom right**), a pattern more consistent with multistep amplification than ecDNA formation. **c**, In NCI-H82, AA classified the *MYC* amplicon as ecDNA (**top left**). The amplicon comprises complex genomic rearrangements in 8q and connections to 14q (**bottom left**). Although the CN transition from non-amplified to amplified region shows a high junctional CN at the pseudobulk level (**bottom right**), the single-cell CN distribution is consistent with ICamp (**top right**). Metaphase fluorescence *in situ* hybridization (FISH) revealed an HSR and no evidence of ecDNA (**Extended Data** Fig. 4), consistent with prior ecDNA amplification followed by chromosomal integration.

## Notes

### Summary of Updates

Figure 1 and 2 revised; Supplemental files updated.

